# Lactate Receptor Activation Alleviates Senescence and Preserves Homeostasis of Aged Arteries

**DOI:** 10.64898/2026.09.20.753049

**Authors:** Yulun Wu, Hamsa Vardini Senthil Kumar, Sai Harsha Bhamidipati, Haibo Yu, Pihu Mehrotra, Pedro Lei, Arundhati Das, Patrick Saari, Maryam Elsayed, Linzhang Huang, Stelios T. Andreadis

## Abstract

Arteries are among the first tissues to exhibit age-related dysfunction, yet the metabolic mechanisms driving vascular senescence remain poorly understood. Here, analysis of human aortic transcriptomic data identified HCAR1, encoding the lactate receptor GPR81, as one of the genes most significantly downregulated with age. We therefore investigated whether age-associated loss of GPR81 contributes to cellular senescence within the vessel wall. Senescent human endothelial cells and vascular smooth muscle cells accumulated neutral and oxidized lipids and exhibited increased labile iron and ferroptosis. Silencing GPR81 in early-passage cells recapitulated this metabolic phenotype together with multiple hallmarks of cellular senescence. Moreover, endothelial-specific deletion of GPR81 in young mice was sufficient to induce senescent cell accumulation, impaired lipid homeostasis, endothelial dysfunction, and elastin disorganization. Conversely, pharmacological activation of GPR81 with the agonist CHBA restored fatty acid metabolism, promoted glycolytic reprogramming, and attenuated ferroptotic stress and senescence-associated phenotypes. In lamin A knock-in (LAKI) progeroid mice, CHBA reduced arterial lipid accumulation and cellular senescence, shifted vascular cell composition toward a youthful state, improved endothelial integrity, and restored extracellular matrix homeostasis. Together, these findings identify age-associated loss of GPR81 as a driver of vascular metabolic dysfunction and cellular senescence and establish pharmacological GPR81 activation as a promising therapeutic strategy for preserving vascular homeostasis and mitigating age-associated cardiovascular disease.

## Introduction

Aging is a complex biological process characterized by progressive decline in cellular and organ function, which increases susceptibility to chronic diseases. Recent human proteomic mapping suggests that arteries are among the earliest and most sensitive tissues to undergo age-dependent remodeling. Moreover, arteries may act as drivers of systemic aging by secreting circulating senescence-associated proteins into plasma^1^. This reinforces Osler’s observation that “Longevity is a vascular question, and man is only as old as his arteries”, emphasizing the centrality of vascular aging in organismal healthspan^2^.

Within the aging vessel wall, senescent cells accumulate. These cells are metabolically active but irreversibly withdraw from the cell cycle. They are resistant to apoptosis and secret senescence-associated secretory phenotype (SASP) factors comprising proinflammatory cytokines, chemokines, growth factors and extracellular matrix (ECM) proteins, which propagate dysfunction to neighboring cells and disrupt tissue homeostasis^3^. In the vasculature, accumulation of senescent vascular cells with age accelerates harmful arterial remodeling, endothelial dysfunction and vascular stiffening, thereby fueling the pathogenesis of age-related cardiovascular diseases (CVDs) including atherosclerosis, stroke, myocardial infarction, and heart failure^4–6^. Given that CVDs remain the leading cause of global mortality, resulting in nearly 0.9–1.0 million deaths per year in the United States^7^, there is an increasing need to identify new therapeutic strategies that target vascular senescence.

Lipid metabolism is a key determinant of vascular aging and CVDs. Genetic studies have identified at least 54 lipid metabolism-related genes causally associated with CVDs, of which 29 represent candidate diagnostic biomarkers or therapeutic targets^8^. Lipids serve not only as structural components of cellular membranes but also act as bioactive signaling molecules, regulating longevity pathways such as insulin/IGF-1 signaling (IIS), mechanistic target of rapamycin (mTOR) signaling, and germline endocrine axes^9^. Senescent vascular cells display profound metabolic reprogramming, including altered fatty acid (FA) utilization and a shift towards anabolic lipid synthesis that fuels SASP production^5,10^. However, whether fatty acid oxidation (FAO) is enhanced or reduced in senescent vascular cells remains controversial. Some studies suggest FAO is enhanced to support the SASP^11–13^, while others report that boosting FAO extends lifespan and impaired FAO accelerates vascular aging^5,14,15^.

Dysfunctional FA metabolism further results in the accumulation of ceramide, lipid droplets and free cytosolic poly unsaturated FAs (PUFAs) due to ineffective lipid utilization, which predispose senescent cells to lipotoxic stress^16–18^. PUFAs are highly susceptible to peroxidation, which generates reactive products that act as lipid-based SASP mediators. Their accumulation promotes oxidative stress, endoplasmic reticulum (ER) stress, mitochondrial dysfunction and cellular senescence^19^. Therefore, restoring lipid homeostasis may be critical to reverse vascular senescence.

Lipid peroxidation is also the initiating event of ferroptosis, an iron-dependent programmed cell death, has recently been implicated in vascular pathology. Accumulating evidence demonstrates that ferroptosis promotes vascular senescence, aneurysm progression and vascular stiffening, thereby exacerbating cardiovascular aging^20–24^. Mechanistically, ferroptosis is tightly coupled to lipid metabolism through lipid-handling enzymes, such as acyl-CoA synthetase long chain family member 4 (ACSL4), lipoxygenases (ALOX15) as well as stearoyl-CoA desaturase 1 (SCD1), which regulate the accumulation and detoxification of lipid peroxides^25–27^. These observations position ferroptosis as a mechanistic link between lipid dysregulation and vascular senescence. Thus, identifying upstream metabolic checkpoints that regulate lipid metabolism to prevent lipid peroxidation and ferroptosis may offer a new approach to reverse vascular senescence, and further slow or reverse vascular aging.

The lactate receptor G protein-coupled receptor 81 (GPR81, also known as hydroxycarboxylic acid receptor 1, HCAR1), has emerged as a role linking metabolic state to lipid handling. GPR81 is a Gi/o coupled G protein-coupled receptor (GPCR) that suppresses lipolysis in adipose tissue by inhibition the cyclic adenosine monophosphate (cAMP) – protein kinase A (PKA) axis, thereby regulating systemic lipid availability^28,29^. Although its lipid regulatory function was originally characterized in adipose tissue, GPR81 expression is also reported in cardiovascular tissues^30^. Recent studies suggest that activation of GPR81 influences on vascular inflammation, endothelial barrier function, and atherosclerotic remodeling, which are tightly linked with vascular senescence^31–33^. However, its specific function in vascular senescence remains undefined.

In this study, because our analysis of human aortic transcriptomes identified HCAR1 as significantly downregulated with age, we hypothesized that GPR81 signaling could maintain lipid homeostasis, thereby mitigating lipid peroxidation and ferroptosis, and ultimately attenuating vascular senescence and restoring vascular integrity. To test this, we employed (i) replicative senescence models of human umbilical vein endothelial cells (HUVECs) and vascular smooth muscle cells (VSMCs), (ii) lentiviral shRNA-mediated GPR81 knockdown of HUVECs and VSMCs, (iii) endothelial-specific GPR81 deletion in young mice and (iv) pharmacological activation using the GPR81 agonist CHBA. We further validated the geroprotective effects of CHBA in vivo using a progeroid mouse model. Collectively, our findings identify GPR81 as a promising therapeutic target for reversing vascular senescence and potentially resetting cardiovascular aging.

## Results

### 1. Vascular aging is associated with lipid dysregulation, ferroptotic stress and loss of GPR81

Several studies reported the crosstalk between lipid metabolism and longevity in diverse species^34–36^. However, the interlink between vascular senescence and dysregulation of lipid metabolism remains insufficiently understood. To address this, we first examined whether vascular senescence is intrinsically associated with lipid dysregulation and ferroptosis. We employed human umbilical vein endothelial cells (HUVECs) and human aortic smooth muscle cells (HASMCs), which represent two of the key vascular cell types, namely endothelial cells (ECs) and vascular smooth muscle cells (VSMCs), respectively. Early-passage cells were used as controls (C), while late-passage cells were used to model replicative senescence (S) **(Fig. 1A)**. The senescence phenotype was first validated by senescence-associated beta-galactosidase (SA-β-gal) enzymatic activity **(Fig. 1B)**, a well-known biomarker of cellular senescence. As expected, S cells of both cell types showed significantly elevated β-gal activity, confirming the senescent phenotype.

**Figure 1:**
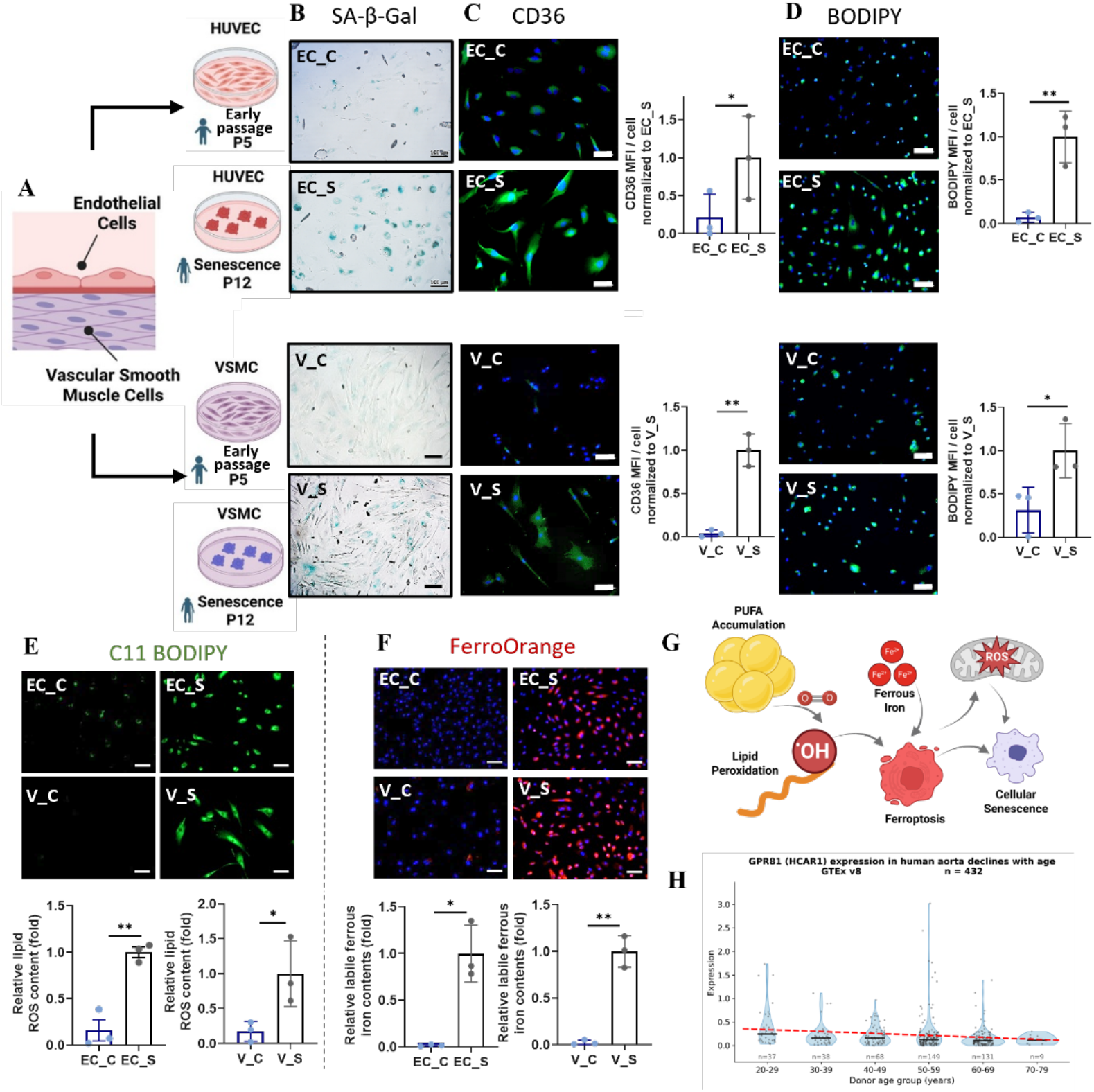
Vascular aging is associated with lipid dysregulation and ferroptotic stress. **A)** Schematic overview of experimental design. **B)** Senescence-associated β-galactosidase (SA-β-Gal) staining in early-passage (EC_C, V_C) and senescent (EC_S, V_S) cells. **C)** Immunofluorescence images and quantification of CD36 expression (green) in C and S cells. Nuclei were counterstained with Hoechst 33342 (blue). Scale bar, 50 µm. Data shown as mean ± SD of n=3 independent experiments, normalized to S. **D)** Representative images and quantification for BODIPY 493/503 (green) staining shows lipid droplets in C and S cells. Nuclei were counterstained with Hoechst 33342 (blue). Scale bar, 100 µm. Data shown as mean ± SD of n=3 independent experiments, normalized to S. **E)** C11-BODIPY fluorescence imaging and quantification to assess oxidized lipid species in C and S cells. Scale bar, 100 µm. Data shown as mean ± SD of n=3 independent experiments, normalized to S. **F)** FerroOrange staining and quantification to visualize intracellular Fe²⁺ levels in C and S cells. Nuclei were counterstained with Hoechst 33342 (blue). Scale bar, 100 µm. Data shown as mean ± SD of n=3 independent experiments, normalized to S. **G)** Schematic illustration depicting the relationship between PUFA accumulation, lipid peroxidation and ferroptotic stress during cellular senescence. **H)** Violin plot of HCAR1 expression in human aorta (GTEx v8, n=432) grouped by donor age. Individual donors are shown as points; black bars indicate group median, and the red dashed line indicates the linear trend across age groups.

Next, we investigated whether vascular senescence is accompanied by altered lipid metabolism. We observed a significant upregulation of CD36 expression, a fatty acid (FA) translocase located on the plasma membrane, in senescent EC (EC_S) and senescent VSMC (V_S) cells, compared to early passage controls **(Fig. 1C)**. Specifically, CD36 expression increased by 4.6-fold (p < 0.05) in EC_S and 26.3-fold (p < 0.01) in V_S, suggesting enhanced lipid uptake in senescent vascular cells. Indeed, BODIPY staining revealed pronounced intracellular lipid accumulation in senescent vascular cells (EC_S: 14.0-fold, p < 0.01; V_S: 3.1-fold, p < 0.05), relative to early passage control cells **(Fig. 1D)**.

Given the increased lipid accumulation, whether this accumulated lipid was also oxidatively damaged remained to be determined. We therefore evaluated lipid peroxidation using the oxidized lipid probe C11-BODIPY. The results showed significant elevation of oxidized lipid probe (green) in senescent vascular cells of both types **(Fig. 1E)**. Specifically, the fluorescence intensity increased by 6.2-fold (p < 0.01) and 5.7-fold (p < 0.05) in senescent EC_S and V_S, respectively, compared to control cells. Lipid peroxidation is a key trigger of ferroptosis, an iron-catalyzed cell death typically associated with age-related diseases^37^, and C11-BODIPY is widely recognized as a ferroptosis-sensitive probe^38^. To investigate whether increased lipid peroxidation caused ferroptotic stress, we detected for the presence of ferrous ion using the ferrous ion binding dye, FerroOrange. The intracellular iron content was significantly elevated in senescent vascular cells (EC_S: 66.6-fold, p < 0.05; V_S: 43.4-fold increase, p < 0.01), as compared to minimal levels in control cells **(Fig. 1F)**. Collectively, these findings demonstrate that replicative senescence in vascular cells is tightly coupled to lipid dysregulation and increased ferroptotic stress **(Fig. 1G)**.

These findings identify impaired lipid handling as a hallmark of vascular senescence, prompting us to investigate the upstream receptor that may be mitigating this response. GPR81 has been linked to lipid metabolism in adipose and skeletal muscle cells. Therefore, we hypothesized that GPR81 may play a critical role in regulating lipid dysregulation and ferroptotic stress in senescent vascular cells. RNA-seq analysis of 432 histologically normal human aortic samples from the Genotype-Tissue Expression (GTEx) v8 project^39^ revealed a significant age-associated decline in *HCAR1*, the gene encoding GPR81 (FDR q=4.9 x 10^-5^) **(Fig. 1H)**. Similarly, analysis of single-cell RNA-sequencing data from mouse aortas in the Tabula Muris Senis dataset^40^, accessed through CZ CELLxGENE Discover^41^, revealed a significant age-associated decline in *Hcar1* expression in 24-month-old compared with 3-month-old mice **(Supplementary Fig. S1A)**. This age-associated decline was further validated at the protein level, as GPR81 expression was significantly reduced in the aortas of both 24-month-old mice and homozygous lamin A knock-in (LAKI) progeroid mice, a model of premature aging, compared with 3-month-old wild-type controls **(Supplementary Fig. S1B)**. LAKI mice harboring the homozygous mutation (LmnaG609G/G609G) exhibit a markedly shortened lifespan of approximately 3–4 months and develop many age-associated phenotypes at an accelerated rate, including cardiovascular abnormalities such as increased arterial stiffness, chronic inflammation, and severe depletion of vascular smooth muscle cells (VSMCs) from the vascular wall^42–44^. Together, these findings identify reduced GPR81 expression as a conserved feature of vascular aging in humans, naturally aged mice, and the LAKI model of premature aging.

### 2. Knockdown of GPR81 leads to lipid dysregulation and enhanced markers of cellular senescence

To further test whether loss of GPR81 contributes to vascular senescence, we knocked down GPR81 in EC_C (EC_Y_shGPR81) and V_C (V_Y_shGPR81) using lentivirus encoded shRNA **(Fig. 2A)**, and GPR81 depletion was confirmed by immunoblotting **(Fig. 2B)**. Early passage control cells transduced with the same shLVDP vector that did not encode any shRNA served as negative controls (EC_Y_vec and V_Y_vec). Strikingly, GPR81 knockdown led to markedly disturbed lipid homeostasis in both cell types. Specifically, BODIPY staining revealed significantly increased lipid accumulation in EC_Y_shGPR81 (48.8-fold, p < 0.01) and V_Y_shGPR81 (13.1-fold, p < 0.05) compared to controls **(Fig. 2C)**. Likewise, mean fluorescence intensity (MFI) of CD36 was elevated in both EC_Y_shGPR81 (97.5-fold, p < 0.01) and V_Y_shGPR81 (372.8-fold, p < 0.01) **(Fig. 2D)**. In addition, following GPR81 knockdown, both cell types exhibited a significant increase in oxidized lipid content (EC_Y_shGPR81: 11.1-fold, p < 0.05; V_Y_shGPR81: 35.5-fold, p < 0.01) **(Fig. 2E)**, as measured by C11 BODIPY. Finally, FerroOrange staining revealed elevated intracellular Fe²⁺ levels in EC_Y_shGPR81 (340.6-fold increase, p < 0.05) and V_Y_shGPR81 (155.8-fold increase, p < 0.05) **(Fig. 2F)**. These results indicate a strong link of GPR81 to lipid metabolism and ferroptotic stress.

**Figure 2:**
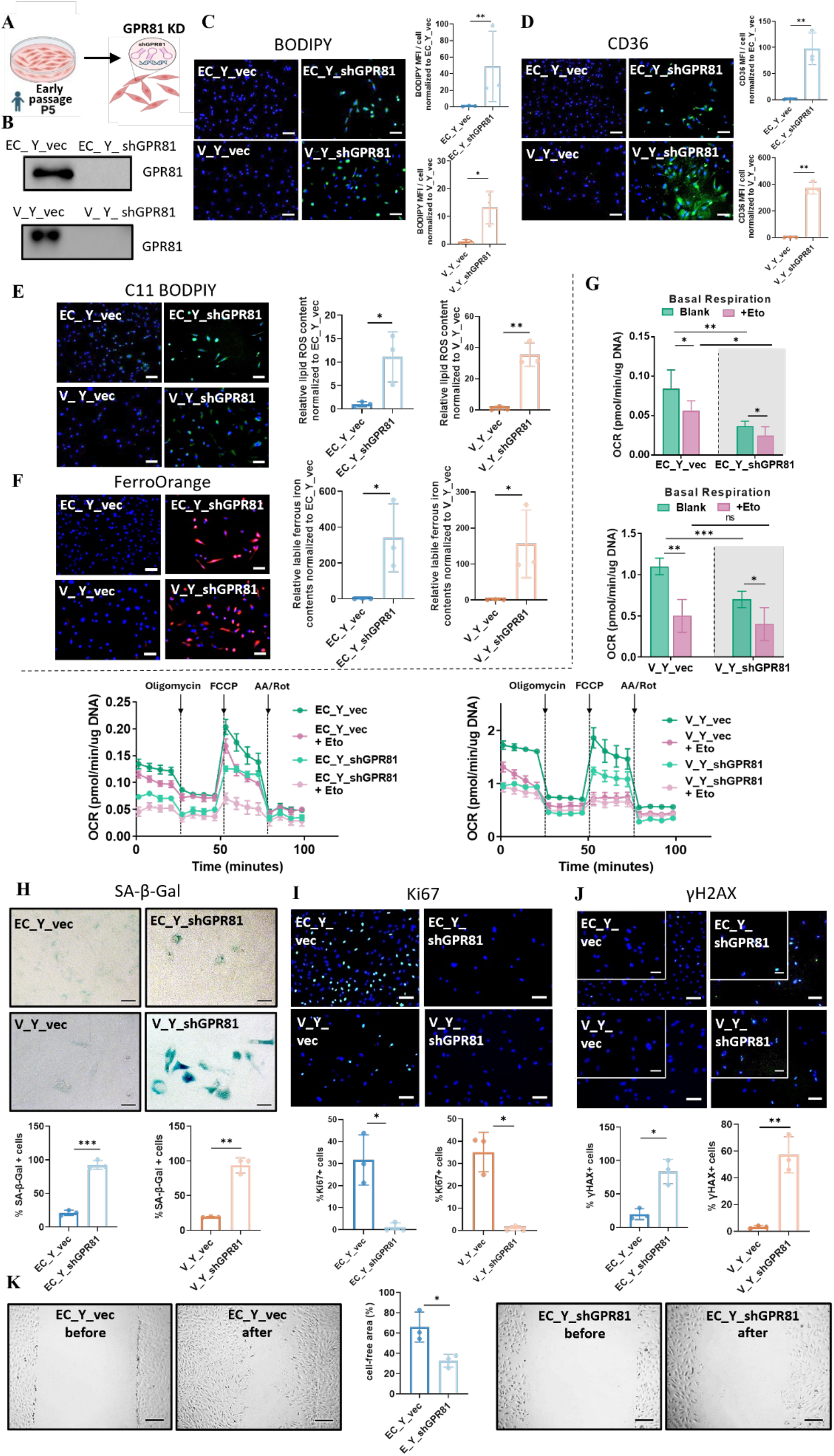
Knockdown of GPR81 leads to lipid dysregulation and enhanced markers of cellular senescence. **A)** Schematic illustrating the experimental approach. GPR81 was silenced in early passage primary HUVECs and VSMCs using shRNA (shGPR81). **B)** Immunoblot analysis of GPR81 expression in vector control (EC_Y_vec and V_Y_vec) and knockdown (EC_Y_shGPR81 and V_Y_shGPR81) ECs and VSMCs. **C)** Representative images and quantification for BODIPY 493/503 (green) staining in control and shGPR81 knockdown ECs and VSMCs to visualize neutral lipid content. Nuclei were counterstained with Hoechst 33342 (blue). Scale bar, 100 µm. Data shown as mean ± SD of n=3 independent experiments, normalized to Y_vec. **D)** Immunofluorescence staining of CD36 in control and shGPR81 knockdown ECs and VSMCs. Nuclei were counterstained with Hoechst 33342 (blue). Scale bar, 100 µm. Data shown as mean ± SD of n=3 independent experiments, normalized to Y_vec. **E)** Representative C11-BODIPY fluorescence imaging and quantifications of oxidized lipid species in control and shGPR81 knockdown ECs and VSMCs. Nuclei were counterstained with Hoechst 33342 (blue). Scale bar, 100 µm. Data shown as mean ± SD of n=3 independent experiments, normalized to Y_vec. **F)** FerroOrange staining used to assess intracellular Fe²⁺ levels in control and shGPR81 knockdown ECs and VSMCs. Nuclei were counterstained with Hoechst 33342 (blue). Scale bar, 100 µm. Data shown as mean ± SD of n=3 independent experiments, normalized to Y_vec. **G)** Seahorse extracellular flux analysis performed to assess FAO. Panels include basal respiration measurements under blank conditions (green) or etomoxir (Eto) treatment (pink) calculated from FAO-OCR data. Data shown as mean ± SD. **H)** SA-β-Gal staining and quantification of % SA-β-Gal+ of control and shGPR81 knockdown ECs and VSMCs. Scale bar, 50 µm. Data shown as mean ± SD of n=3 independent experiments. **I)** Immunofluorescence staining of Ki67 and quantification of % Ki67+ cells. Nuclei were counterstained with Hoechst 33342 (blue). Scale bar, 100 µm. Data shown as mean ± SD of n=3 independent experiments. **J)** Immunofluorescence staining for γH2AX in control and shGPR81 knockdown ECs and VSMCs, with quantification of % γH2AX+ cells. Nuclei were counterstained with Hoechst 33342 (blue). Scale bar, 100 µm (bottom) and 50 µm (top). Data shown as mean ± SD of n=3 independent experiments. **K)** Scratch wound–healing assay evaluating cell migration in control and shGPR81 knockdown ECs at 0 and 12 hrs post-wounding, with quantification of cell-free area percentage. Data shown as mean ± SD of n=3 independent experiments.

Since excessive lipid accumulation may reflect defective mitochondrial FAO capacity, we evaluated FAO processes under condition of L-carnitine with substrate-limited medium, based on the oxygen consumption rate (OCR) using a Seahorse XF analyzer **(Fig. 2G)**. After knocking down GPR81, both EC_Y_shGPR81 and V_Y_shGPR81 showed significant reduction in basal respiration compared to control groups (EC_Y_shGPR81: 0.037 ± 0.006 pmol/min/µg DNA, EC_Y_vec: 0.084 ± 0.024 pmol/min/µg DNA, p<0.01; V_Y_shGPR81: 0.7 ± 0.1 pmol/min/µg DNA, V_Y_vec: 1.1 ± 0.1 pmol/min/µg DNA, p<0.001), indicating reduced mitochondrial respiration.

Next, we examined the mitochondrial FAO capacity specifically, by treating cells with etomoxir (Eto), a chemical inhibitor of carnitine palmitoyl transferase 1 (CPT1), which blocks mitochondria import of long-chain FAs (LCFAs) for further β-oxidation **(Fig. 2G)**. The difference in FAO-OCR between etomoxir-treated (+Eto) and untreated (Blank) cells reflects the utilization level of endogenous free fatty acids (FFAs) during β-oxidation. Etomoxir-treated groups (pink) showed reduced basal respiration compared to non-treated groups (green) in both control (EC_Y_vec: 0.056 ± 0.013 pmol/min/µg DNA, p < 0.05; V_Y_vec: 0.5 ± 0.2 pmol/min/µg DNA, p < 0.01) and GPR81-knockdown cells (EC_Y_shGPR81: 0.027 ± 0.011 pmol/min/µg DNA, p < 0.05; V_Y_shGPR81: 0.4 ± 0.2 pmol/min/µg DNA, p < 0.05). The percent decrease was less pronounced in the knockdown groups (EC_Y_shGPR81: 27.0%; V_Y_shGPR81: 42.8%) than in the corresponding control groups (EC_Y_vec: 33.3%, V_Y_vec: 54.5%), indicating that loss of GPR81 compromised endogenous FA utilization, possibly by impairing mitochondrial β-oxidation. Together, these findings highlight GPR81 is essential for maintaining lipid hemostasis in vascular cells, protecting cells from lipid toxicity and further ferroptotic stress.

Because vascular lipid dysregulation and ferroptosis are tightly linked with cellular senescence, we next evaluated the impact of GPR81 deficiency on vascular senescence. Indeed, SA-β-Gal staining revealed elevated senescence in EC_Y_shGPR81 (4.3-fold, p < 0.001) and V_Y_shGPR81 (4.8-fold, p < 0.01) compared to controls **(Fig. 2H)**. In addition, the percentage of Ki67+ (proliferating) cells decreased dramatically in both EC_Y_shGPR81 (from 31.6 ± 11.3% to 1.1 ± 1.9%, p < 0.05) and V_Y_shGPR81 (from 35.1 ± 8.8% to 0.6 ± 1.1%, p < 0.05) **(Fig. 2I)**. Interestingly, a wound healing assay showed that the migration ability of EC was also significantly impaired after GPR81 knock down **(Fig. 2K)**, decreasing from 65.8 ± 14.8% in EC_Y_vec to 32.6 ± 6.3% in EC_Y_shGPR81 (p < 0.05). We also assessed the percentage of DNA damaged cells by staining for γH2AX, an early cellular response to induce DNA double-strand breaks. Compared with EC_Y_vec and V_Y_vec, γH2AX+ cells were significantly higher in both EC_Y_shGPR81 and V_Y_shGPR81 (4.2-fold and 19.0-fold, respectively) **(Fig. 2J)**, indicating increased DNA damage in the absence of GPR81. Overall, these findings show that GPR81 knockdown induces multiple hallmarks of cellular senescence.

Given that senescence is often accompanied by metabolic reprogramming and mitochondrial dysfunction, we further investigated how GPR81 knockdown impairs mitochondrial-dependent energy metabolism, by measuring OCR **(Supplementary Fig. S2A)** and the extracellular acidification rate (ECAR) **(Supplementary Fig. S2B)** using Seahorse XF96 Extracellular Flux analyzer. Although there is no significant difference in basal respiration between V_Y_shGPR81 and V_Y_vec groups, basal respiration was significantly reduced in EC_Y_shGPR81 (0.2 ± 0.2 pmol/min/µg DNA, p < 0.01) compared to EC_Y_vec (0.7 ± 0.2 pmol/min/µg DNA), indicating impaired basal mitochondrial function. Importantly, maximum respiration was significantly reduced in both GPR81-deficient cell types (EC_Y_shGPR81: 0.4 ± 0.1 pmol/min/µg DNA, p < 0.01; V_Y_shGPR81: 1.0 ± 0.4 pmol/min/µg DNA, p < 0.05) compared to control groups (EC_Y_vec: 1.5 ± 0.5 pmol/min/µg DNA; V_Y_vec: 2.0 ± 0.8 pmol/min/µg DNA), indicating a diminished capacity of cells to meet their bioenergetic demands after knocking down GPR81.

Furthermore, ECAR measurements revealed that glycolytic function, including glycolysis and glycolytic capacity, was significantly impaired in EC_Y_shGPR81 (glycolysis: 0.2 ± 0.1 mpH/min/µg DNA, p < 0.01; glycolytic capacity: 0.2 ± 0.1 mpH/min/µg DNA, p < 0.01) and V_Y_shGPR81 (glycolysis: 0.1 ± 0.01 mpH/min/µg DNA, p < 0.05; glycolytic capacity: 0.1 ± 0.01 mpH/min/µg DNA, p < 0.05) relative to controls (EC_Y_vec: glycolysis: 0.9 ± 0.3 mpH/min/µg DNA, glycolytic capacity: 0.9 ± 0.3 mpH/min/µg DNA; V_Y_vec: glycolysis: 0.3 ± 0.1 mpH/min/µg DNA, p < 0.01; glycolytic capacity: 0.3 ± 0.1 mpH/min/µg DNA, p < 0.01). These findings demonstrate that GPR81 loss induces a senescence-like phenotype in vascular cells, characterized by impaired proliferation, increased DNA damage, and compromised mitochondrial oxidative metabolism and glycolytic function.

### 3. Endothelial GPR81 deficiency disturbs vascular development in young mice

Because GPR81 deficiency induced cellular senescence and impaired mitochondrial function, we next investigated whether GPR81 is required to maintain vascular homeostasis in vivo. To this end, we employed a transgenic mouse model that we developed in which GPR81 (Hcar1) is selectively deleted in endothelial cells (ECs) (Hcar1^fl/fl Cdh5-Cre) **(Fig. 3A)**. Firstly, immunostaining for p21 revealed a significantly higher percentage of p21+ luminal ECs in Hcar1^fl/fl^ Cdh5-Cre (3.9-fold increase, p<0.05), compared with control mice (Hcar1^fl/fl^), indicating that endothelial GPR81 deficiency is sufficient to induce senescence **(Fig. 3B)**. Notably, the percentage of p21+ cells was also significantly increased within the medial layer (4.8-fold increase, p<0.05), suggesting that endothelial GPR81 deficiency may have a broader effect in the vascular wall. Moreover, given the link between GPR81 signaling, lipid metabolism and cellular senescence, we assessed CD36 expression in the aortic wall. CD36 expression showed an increasing trend in Hcar1^fl/fl^ Cdh5-Cre mice compared with Hcar1^fl/fl^ controls (1.8-fold increase, p=0.058) **(Fig. 3C)**, suggesting that endothelial GPR81 deficiency may be associated with enhanced lipid uptake.

**Figure 3:**
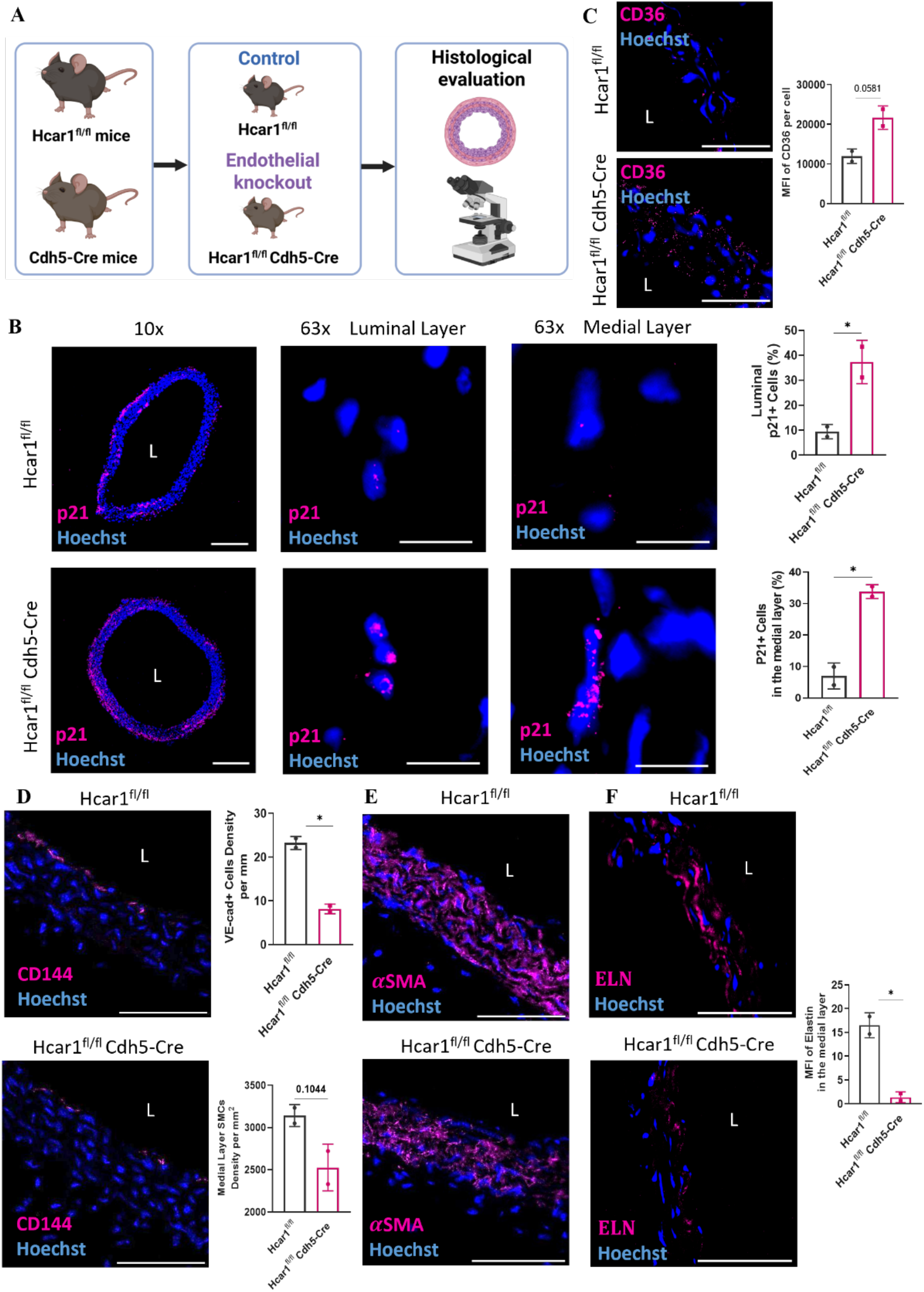
Endothelial GPR81 deficiency disturbs vascular development in young mice. **A)** Schematic showing EC-specific knockdown of GPR81 experimental design using Hcar1^fl/fl^ mice and Hcar1^fl/fl^ Cdh5-Cre mice. **B)** Representative immunofluorescence images and quantitative analysis of % p21+ cells in arteries. L represents lumen. Scale bar, 200 µm (10x) and 10 µm (63x). n = 2 independent biological samples. Data shown as mean ± SD. **C-F)** Representative immunofluorescence images and quantitation of **(C)** CD36, **(D)** CD144, **(E)** *α*SMA and **(F**) Elastin (ELN) in arteries from mice. L represents lumen. Scale bar, 50 µm. n = 2 independent biological samples. Data shown as mean ± SD.

We next asked whether endothelial GPR81 deficiency compromises vascular integrity. Immunostaining for VE-cadherin (CD144), a marker of endothelial junctional integrity, revealed an approximately 65% reduction in Hcar1^fl/fl^ Cdh5-Cre mice (8.1 VE-cadherin+ cells per mm), compared with Hcar1^fl/fl^ mice (23.2 VE-cadherin+ cells per mm, p<0.05), indicating that EC-specific deletion of GPR81 compromises endothelial junctional integrity, even in young mice **(Fig. 3D)**. Interestingly, the density of αSMA+ cells in the medial layer showed a decreasing trend that did not reach statistical significance (p=0.1044) in Hcar1^fl/fl^ Cdh5-Cre mice **(Fig. 3E)**. In addition, elastin (ELN) immunofluorescence, a key structural ECM protein that degrades with age^45^, demonstrated a marked reduction in MFI in the aortic wall of Hcar1^fl/fl^ Cdh5-Cre mice (p<0.05) **(Fig. 3F)**, suggesting impaired elastin organization in young mice following endothelial GPR81 deletion. Overall, our findings suggest that loss of endothelial GPR81 is associated with cellular senescence, altered lipid handling, impaired endothelium integrity and disrupted ECM protein organization in the vascular wall.

### 4. CHBA treatment restores lipid metabolism in senescent vascular cells in vitro

We next investigated whether pharmacological activation of GPR81 could restore metabolic balance and alleviate senescent phenotypes in vascular cells. To achieve this, senescent ECs and VSMCs were treated with the GPR81 agonist^46^, 3-chloro-5-hydroxy BA (CHBA, 100 µM) for 7 days in vitro, and changes in lipid metabolism were assessed. First, CHBA-treated senescent ECs (EC_S+C) and VSMCs (V_S+C) exhibited approximately 4-fold and 5-fold increases in GPR81 mRNA expression, respectively, compared with their untreated senescent counterparts (EC_S and V_S). These findings suggest the existence of a positive feedback mechanism whereby GPR81 activation enhances its own gene expression **(Fig. 4A)**.

**Figure 4:**
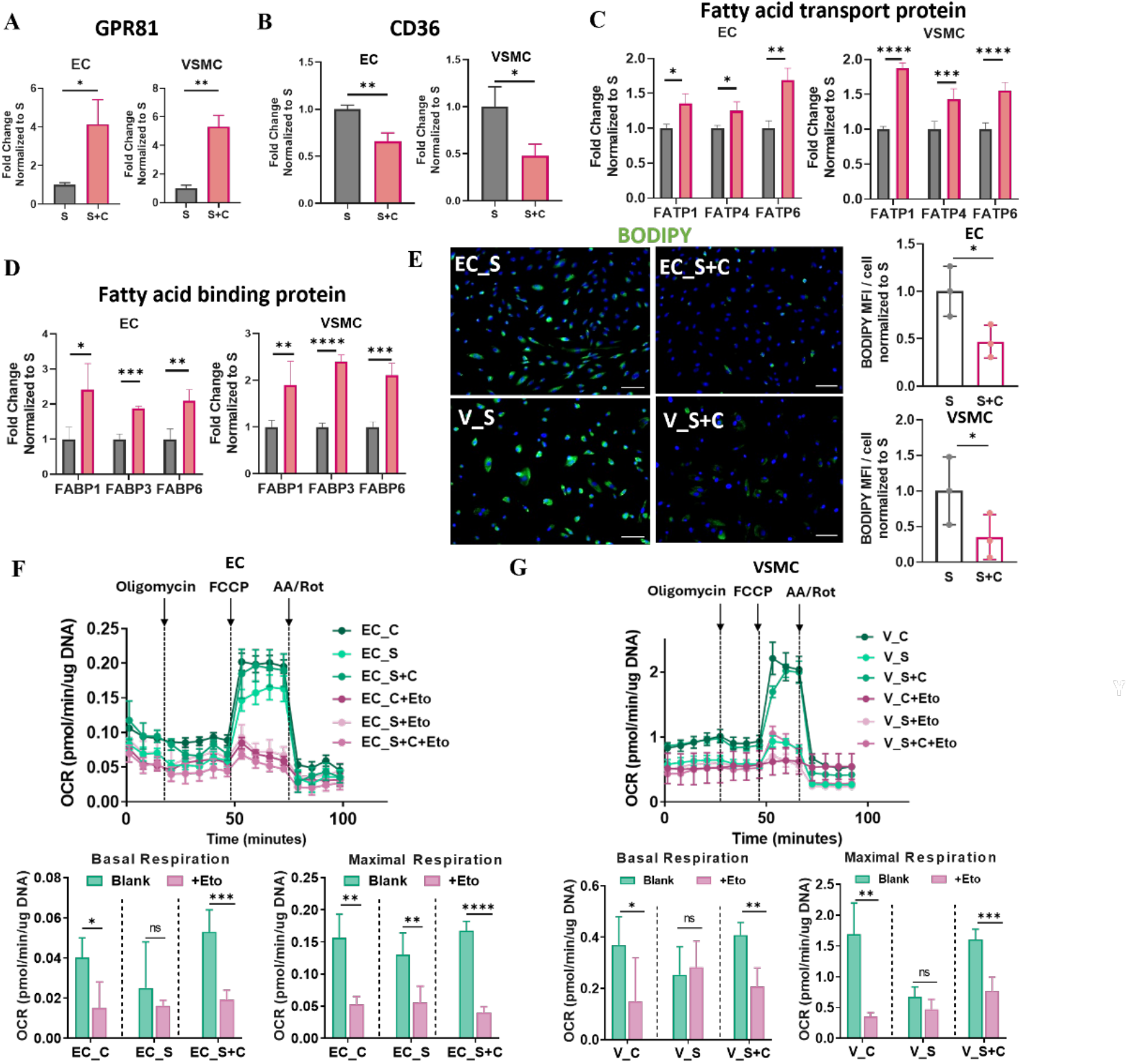
CHBA treatment restores lipid metabolism in senescence vascular cells in vitro. Quantitative RT-PCR for **A)** GPR81, **B)** cytoplasmic fatty acid transport protein CD36, **C)** Fatty Acid Transport Proteins: FATP1, FATP4 and FATP6, and **D)** Fatty Acid Binding Proteins: FABP1, FABP3 and FABP6. Data shown as mean ± SD. **E)** Representative images for BODIPY 493/503 (green) staining depicts lipid droplets in senescent ECs and VSMCs cultured without (S) or with CHBA (S+C), and MFI quantification. Nuclei were counterstained with Hoechst 33342 (blue). Scale bar, 100 µm. Data shown as mean ± SD of n=3 independent experiments, normalized to S. **F-G)** Seahorse extracellular flux analysis of mitochondrial respiration to measure FAO in early-passage control (C), senescent (S) and CHBA treated senescent (S+C) **(F)** ECs or **(G)** VSMCs in the absence (green) or presence of etomoxir (Eto) treatment (pink). Basal and maximal respiration values were calculated from FAO-OCR and data are shown as mean ± SD.

Furthermore, we examined whether GPR81 activation affects lipid processing and transport. CHBA treatment downregulated CD36 in senescent cells **(Fig. 4B)**, and upregulated genes involved in fatty acid (FA) import and intracellular trafficking^10^, such as FA transport proteins (FATPs) - *FATP1, FATP4*, and *FATP6* **(Fig. 4C)**. Similarly, FA binding proteins (*FABP1, FABP3,* and *FABP6*), which facilitate intracellular transport of FAs to organelles for oxidation, esterification, or signaling^47^, were also upregulated following CHBA treatment **(Fig. 4D)**. Collectively, these changes suggest that CHBA-mediated GPR81 activation may enhance lipid metabolism including FA uptake and trafficking in senescent vascular cells.

To assess whether these transcriptional changes translated into functional improvements in lipid metabolism, we performed BODIPY staining to visualize intracellular lipid accumulation **(Fig. 4E)**. Consistent with these transcriptional changes, a significant reduction in lipid droplet content was observed in both EC_S+C (by 46%, p < 0.05) and V_S+C (by 65%, p < 0.05), demonstrating improved lipid clearance in senescent cells.

Enhanced lipid clearance may suggest augmented mitochondrial FAO, thereby prompting us to evaluate mitochondrial lipid metabolism by measuring FAO-OCR in the presence of etomoxir and carnitine. The difference in FAO-OCR between etomoxir-treated (+Eto) and untreated (Blank) cells reflects the utilization level of endogenous free (F)FAs during β-oxidation. Indeed, the presence of Eto, EC_C exhibited the higher decrease in basal and maximal respiration **(Fig. 4F)** (62% decrease in basal: from 0.04 ± 0.010 pmol/min/µg DNA to 0.015 ± 0.013 pmol/min/µg DNA, p < 0.05; 66% decrease in maximum: from 0.156 ± 0.037 pmol/min/µg DNA to 0.053 ± 0.012 pmol/min/µg DNA, p < 0.01) as compared to EC_S (36% decrease in basal: from 0.025 ± 0.023 pmol/min/µg DNA to 0.016 ± 0.002 pmol/min/µg DNA, p > 0.05; 61% decrease in maximal: from 0.130 ± 0.034 pmol/min/µg DNA to 0.056 ± 0.025 pmol/min/µg DNA, p < 0.01). This result shows that endogenous FAO was impaired in EC_S. Notably, CHBA treatment increased both basal and maximal respiration. Moreover, CHBA-treated senescent ECs (EC_S+C) exhibited a greater reduction in basal and maximal respiration following Eto treatment (64% decrease in basal: from 0.053 ± 0.011 pmol/min/µg DNA to 0.019 ± 0.005 pmol/min/µg DNA, p < 0.001; 76% decrease in maximal: from 0.167 ± 0.015 pmol/min/µg DNA to 0.040 ± 0.009 pmol/min/µg DNA, p < 0.0001), compared with untreated EC_S, indicating enhanced utilization of endogenous fatty acids and restoration of FAO capacity.

Similar trend was observed in senescent VSMCs **(Fig. 4G)**. Specifically, while Eto treatment decreased basal and maximal respiration in V_C (43% decrease in basal: from 0.370 ± 0.110 pmol/min/µg DNA to 0.208 ± 0.072 pmol/min/µg DNA, p < 0.05; 79% decrease in maximal: from 1.688 ± 0.510 pmol/min/µg DNA to 0.350 ± 0.067 pmol/min/µg DNA, p < 0.01), it had no significant effect in V_S (basal: from 0.254 ± 0.109 pmol/min/µg DNA to 0.283 ± 0.102 pmol/min/µg DNA, p > 0.05; maximal: from 0.671 ± 0.163 pmol/min/µg DNA to 0.470 ± 0.163 pmol/min/µg DNA, p > 0.05), suggesting that V_S had lost their ability to utilize endogenous FAs as an energy source. On the other hand, V_S+C exhibited elevated basal and maximal FAO-OCR compared to untreated V_S, and Eto decreased both basal and maximal respiration to similar levels as V_C (49% decrease in basal: from 0.408 ± 0.050 pmol/min/µg DNA to 0.208 ± 0.072 pmol/min/µg DNA, p < 0.01; 52% decrease in maximal: from 1.606 ± 0.168 pmol/min/µg DNA to 0.768 ± 0.227 pmol/min/µg DNA, p < 0.001), indicating that CHBA restored mitochondrial utilization of endogenous FAs.

Taking together, these results demonstrate that GPR81 activation by CHBA restores lipid metabolism as evidenced by clearing excess lipid accumulation, enhancing lipid trafficking genes and partially restoring endogenous FAO.

### 5. CHBA treatment suppresses age-related ferroptotic stress and senescence-associated phenotype in vascular cells in vitro

Because CHBA restored lipid homeostasis and β-oxidation in senescent vascular cells, we next asked whether these improvements might also mitigate ferroptotic stress, a cell-death program tightly coupled to lipid dysregulation. To explore this, we first examined the expression of key ferroptosis regulators. At the transcriptional level, CHBA treatment significantly suppressed two key drivers of ferroptosis, *ACSL4* and *TFR1* (transferrin receptor protein 1), in both cell types. ACSL4 promotes lipid peroxidation, whereas TFR1 facilitates iron uptake^48–50^ **(Fig. 5A-B)**. To examine whether these transcriptional changes translated into functional outcomes, we assessed lipid peroxidation and intracellular iron accumulation. C11-BODIPY staining revealed that CHBA significantly reduced oxidized lipid levels by 34% in EC_S (p < 0.05) and 70% in V_S (p < 0.05), compared to non-treated senescent cells **(Fig. 5C)**. Reactive oxygen species (ROS) levels were also diminished following CHBA treatment **(Supplementary Fig. S3A)**. In parallel intracellular ferrous iron levels **(Fig. 5D)**, assessed by FerroOrange staining, were also markedly reduced by 81% in EC_S (p < 0.01) and 74% in V_S (p < 0.05) following CHBA treatment. Together, these results indicate that CHBA treatment effectively alleviated ferroptotic stress in senescent vascular cells.

**Figure 5:**
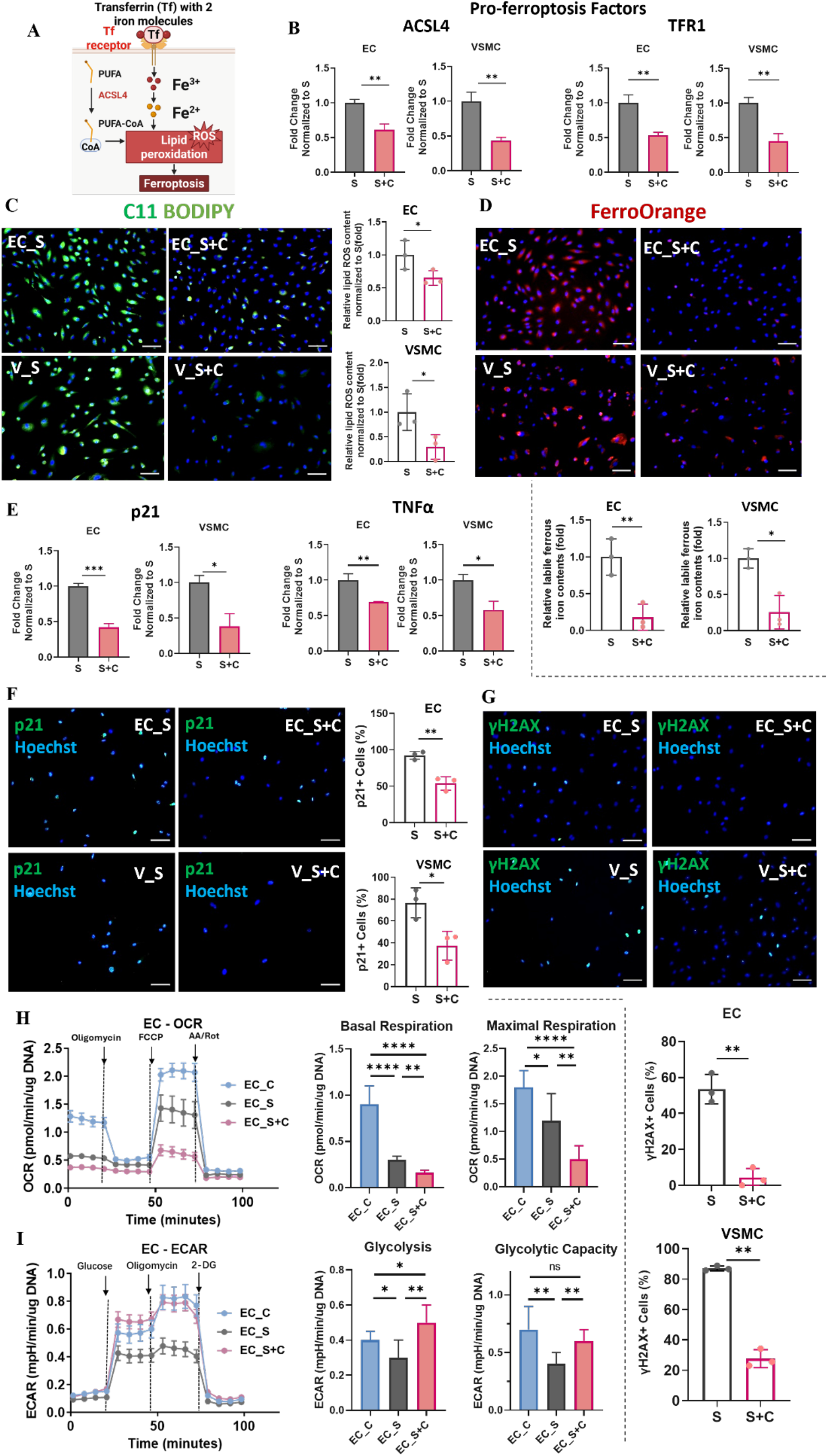
CHBA treatment suppresses age-related ferroptotic stress and senescence-associated hallmarks in vascular cells in vitro. **A)** Schematic showing the ferroptosis regulation pathway. **B)** Quantitative RT-PCR for pro-ferroptosis factors: ACSL4 and TFR1 in the absence (S) or presence of CHBA (S+C). Data were internally normalized to the respective expression level of RPL32 and then normalized to S; data shown as mean ± SD. **C)** Lipid peroxidation in senescent ECs or VSMCs was assessed using C11-BODIPY live staining (green) in the S and S+C; and corresponding quantifications. Nuclei were counterstained with Hoechst 33342 (blue). Scale bar, 100 µm. Data shown as mean ± SD of n=3 independent experiments, normalized to S. **D)** Intracellular ferrous iron pool was measured using FerroOrange probe (red) in S and S+C ECs (top row) and VSMCs (bottom row). Scale bar, 100 µm. Data shown as mean ± SD of n=3 independent experiments, normalized to S. **E)** Quantitative RT-PCR for senescence marker, p21 and TNFα. Data were internally normalized to the respective expression level of RPL32 and then normalized to S; data shown as mean ± SD. **F-G)** Immunostaining for **(F)** p21 (green) and **(G)** γH2AX (green) in S and S+C for both ECs and VSMCs, and corresponding quantifications. Nuclei were counterstained with Hoechst 33342 (blue). Scale bar, 100 µm. Data shown as mean ± SD of n=3 independent experiments. **H)** Seahorse extracellular flux analysis of mitochondrial respiration in EC. Oxygen consumption rate (OCR) following sequential injections (oligomycin, FCCP, antimycin A/rotenone) and quantifications of basal and maximal respiration. Data shown as mean ± SD. **I)** Extracellular acidification rate (ECAR) of EC following sequential injections (glucose, oligomycin, and 2-DG) and quantification of glycolysis and glycolytic capacity. Data shown as mean ± SD.

Given the interlink between ferroptotic stress, lipid metabolism, and vascular senescence, we evaluated whether CHBA could also reverse broader senescence-associated phenotypes in vascular cells. Indeed, the canonical cell cycle arrest markers p21 and p53 were significantly downregulated upon CHBA treatment in both EC_S and V_S **(Fig. 5E, Supplementary Fig. S3B)**. In addition, CHBA treatment markedly downregulated the SASP^51,52^, as indicated by reduced expression of inflammatory cytokines and adhesion molecules, including tumor necrosis factor alpha (TNF-α) **(Fig. 5E)**, intercellular adhesion molecule 1 (ICAM-1), and monocyte chemoattractant protein-1 (MCP-1) **(Supplementary Fig. S3C)**. Consistent with gene expression data, immunofluorescence staining showed a significant reduction in the number of p21⁺ and γH2AX⁺ cells in both EC_S (p21: decreased by 41%, p < 0.01; γH2AX: decreased by 92%, p < 0.01) and V_S (p21: decreased by 51%, p < 0.05; γH2AX: decreased by 68%, p < 0.01) following CHBA treatment **(Fig. 5F-G)**. Collectively, these findings demonstrate that CHBA effectively alleviated inflammation and cellular senescence, reversing several hallmarks of vascular cell senescence.

Given that vascular senescence is closely associated with metabolic reprogramming and mitochondrial dysfunction, we further investigated whether CHBA affects mitochondrial metabolism in vascular senescent cells by measuring OCR and ECAR using Seahorse XF96 Extracellular Flux analyzer. Compared with early-passage EC_C, senescence significantly reduced basal respiration from 0.90 ± 0.20 pmol/min/µg DNA to 0.30 ± 0.03 pmol/min/µg DNA (p < 0.0001), and maximal respiration from 1.80 ± 0.30 pmol/min/µg DNA to 1.19 ± 0.48 pmol/min/µg DNA (p < 0.05) **(Fig. 5H)**. CHBA treatment further suppressed oxidative phosphorylation, decreasing both basal respiration (0.16 ± 0.02 pmol/min/µg DNA, p < 0.01) and maximal respiration (0.49 ± 0.24 pmol/min/µg DNA, p < 0.01) significantly compared with untreated EC_S. A similar trend was observed in VSMCs **(Supplementary Fig. S3D)**. V_S exhibited modestly reduced basal (from 2.2 ± 0.3 to 1.9 ± 0.3 pmol/min/µg DNA, p > 0.05) but significantly impaired maximal respiration (from 8.9 ± 1.5 to 5.7 ± 1.1 pmol/min/µg DNA, p < 0.001). CHBA treatment did not significantly alter basal respiration but further reduced maximal respiration to 3.6 ± 0.8 pmol/min/µg DNA (p < 0.01 compared to V_S).

To further evaluate the effect of CHBA on glycolytic function by measuring ECAR. EC_S exhibited reduced glycolytic activity **(Fig. 5I)**, decreasing glycolysis from 0.4 ± 0.05 to 0.3 ± 0.1 mpH/min/µg DNA (p < 0.05), and glycolytic capacity from 0.7 ± 0.2 to 0.4 ± 0.1 mpH/min/µg DNA (p < 0.01). CHBA markedly increased glycolysis (0.5 ± 0.1 mpH/min/µg DNA, p < 0.01), and glycolytic capacity (0.6 ± 0.1 mpH/min/µg DNA, p < 0.01) to levels comparable to EC_C (glycolysis: p < 0.05; glycolytic capacity: p > 0.05). V_S exhibited a similar metabolic pattern, exhibiting profoundly suppressed glycolysis (0.4 ± 0.1 to 0.05 ± 0.01 mpH/min/µg DNA, p < 0.0001) and glycolytic capacity (0.4 ± 0.1 to 0.15 ± 0.04 mpH/min/µg DNA, p < 0.0001), both of which were moderately increased by CHBA treatment (glycolysis: 0.08 ± 0.01 mpH/min/µg DNA, p < 0.001; glycolytic capacity: 0.23 ± 0.02 mpH/min/µg DNA, p < 0.001) **(Supplementary Fig. S3E)**. Collectively, these findings indicate that CHBA decreases OCR and increases anaerobic glycolysis in ECs and to a lesser extent in VSMCs, possibly as a way to modulate the cell proliferation and survival.

### 6. Single-nucleus RNA sequencing reveals vascular remodeling following CHBA treatment

After demonstrating the senotherapeutic ability of CHBA to restore lipid utilization and reverse senescence hallmarks in vascular cells in vitro, we examined whether CHBA has geroprotective effects in vivo using LAKI mouse, a model of premature aging **(Fig. 6A)**. Homozygous LAKI mice received intraperitoneal (I.P.) injections of CHBA (10 mg/kg), three injections per week, beginning at 10 weeks of age and continuing for four weeks. Mice that received only the vehicle (DMSO in corn oil) served as controls. Blood metabolic panel analysis at the end of the treatment period showed that serum glucose levels were significantly reduced in CHBA-treated mice compared with their pre-treatment levels (p < 0.01), whereas glucose levels in untreated LAKI mice remained unchanged **(Fig. 6B)**.

**Figure 6:**
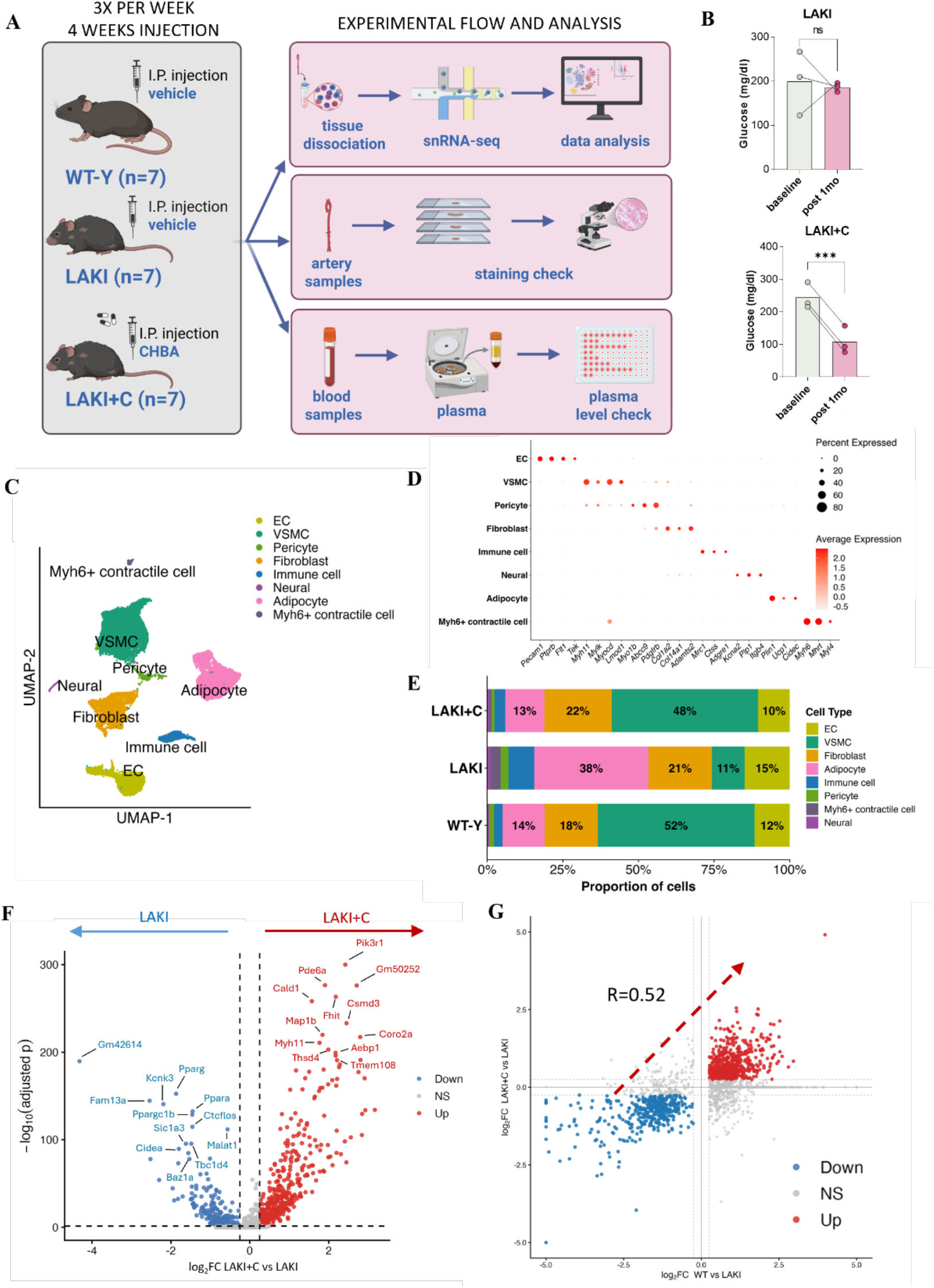
Single-nucleus sequencing investigating CHBA-mediated vascular remodeling in LAKI mice. **A)** Schematic showing experimental timeline; ten week old LAKI mice were administered either vehicle or CHBA intraperitoneally. The arteries were harvested after 4 weeks for further assessments. **B)** Serum blood glucose before (baseline) and after (post 1 month) intraperitoneal injections of CHBA in LAKI and LAKI+C mice. **C)** UMAP visualization of scRNA-seq data shows 8 cell clusters. **D)** Dot plot shows the expression of representative maker genes used for cell type annotation across the identified clusters. **E)** Stacked bar plot showing the proportion of each cell type in young WT, LAKI and LAKI+C arteries. **F)** Volcano plot of differentially expressed genes (DEGs) in CHBA-treated arteries compared with control LAKI arteries. Upregulated DEGs are shown in red (log2FC > 0.25 and P adj. <0.05), downregulated DEGs are shown in blue (log2FC <−0.25 and P adj. <0.05). Top 12 up- and 12 down-regulated DEGs are labeled. Unchanged genes are depicted in gray. **G)** Rescued scatter plot comparing changes in LAKI+C vs. LAKI and WT-Y vs. LAKI. Red dots represent upregulated DEGs in the same direction; blue dots represent downregulated DEGs in the same direction for LAKI+C vs. LAKI and WT-Y vs. LAKI.

To further define how CHBA treatment alters the arterial transcription, we performed single-nucleus RNA sequencing (snRNA-seq) on arteries from young C57BL/6 J (WT-Y, 3 month old, n=7) mice, LAKI mice (n=7) and CHBA treated LAKI mice (LAKI+C, n=7), capturing approximately 29,197 nuclei (314 genes per nuclei). After stringent quality control, Seurat-based clustering identified 8 major cell clusters **(Fig. 6C)**, which were manually annotated based on the expression of established canonical marker genes **(Fig. 6D)**, and further supported by feature plot visualization of cluster defining markers^53–56^ **(Supplementary Fig. S4A-H)**. Neural, Adipocyte and Myh6+ contractile cell clusters were likely attributable to sample contamination from adjacent perivascular or cardiac tissues during sample preparation. Intriguingly, comparison of cell-type composition across groups revealed that CHBA-treated arteries exhibited an increased proportion of VSMCs and a decreased proportion of adipocytes compared with LAKI arteries **(Fig. 6E)**. Notably, their cellular composition closely resembled that of WT-Y arteries.

Next, we compared the differentially expressed genes (DEGs) to evaluate transcriptomic signatures following CHBA treatment. Differential expression analysis identified 396 upregulated and 184 downregulated DEGs in LAKI+C relative to LAKI, indicating transcriptional reprogramming induced by CHBA treatment. Among the top 12 DEGs, genes related to smooth muscle contraction (Cald1, Myh11) and ECM organization (Thsd4, Aebp1) were upregulated in LAKI+C aortas, whereas lipid responsive genes (Pparg, Ppargc1b, Ppara, Cidea) were downregulated compared to LAKI aortas **(Fig. 6F)**. To further evaluate whether these transcriptomic changes reflected a shift toward a more youthful transcriptional state, we compared DEGs identified in LAKI+C vs. LAKI with those identified in WT-Y vs. LAKI arteries. Rescue scatter plot revealed a positive correlation (R=0.52) **(Fig. 6G)**, supporting a partial restoration of the LAKI arterial transcriptome toward the WT-Y transcriptional profile following CHBA treatment.

### 7. Administration of CHBA alleviates vascular lipid accumulation and senescence in vivo

To investigate the functional relevance of the aging-associated transcriptional changes following CHBA treatment, we identified genes that were upregulated in the progeroid vasculature of LAKI mice relative to age-matched WT-Y controls and subsequently downregulated by CHBA treatment. To this end, we generated a ‘recovered-down’ gene set comprising the intersection of DEGs that were downregulated in LAKI+C vs. LAKI (n=184) and upregulated in LAKI vs. WT-Y (equivalently, downregulated in WT-Y vs. LAKI; n=257) **(Fig. 7A)**. Gene Ontology (GO) analysis of the overlapping genes (n=108) revealed significant enrichment of pathways related to lipid metabolism, including “lipid/cholesterol storage”, “triglyceride biosynthetic process”, “carboxylic acid biosynthetic process”, “fatty acid biosynthetic process”, and the “peroxisome proliferator-activated receptor (PPAR) signaling pathway” **(Fig. 7B)**.

**Figure 7:**
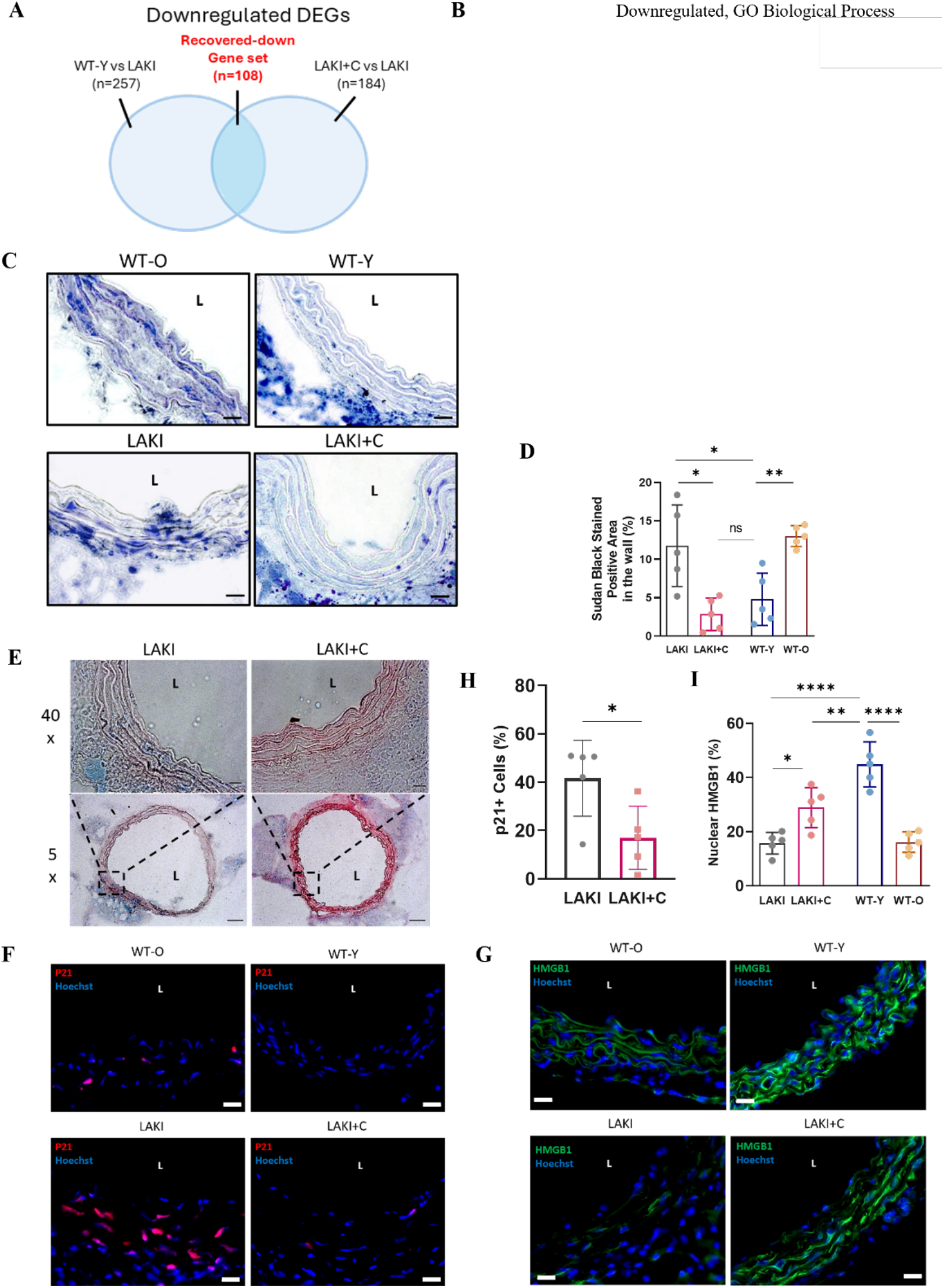
Administration of CHBA alleviates vascular lipid accumulation and senescence in LAKI mice. **A)** Venn diagram of the recovered-down gene sets. **B)** GO enrichment analysis of recovered-down gene sets, highlighting Biological Processes associated with fatty acids. Bubble size represents gene count. **C)** Representative Sudan Black B staining images in arteries from four groups of mice: old WT (WT-O), WT-Y, LAKI and LAKI+C mice. L represents lumen. Scale bar represents 20 µm. **D)** Quantification of the % area that is positive for Sudan Black B, n=5 biological replicates. Data shown as mean ± SD. **E)** Representative SA-β-gal staining in arteries from LAKI and LAKI+C mice. L represents lumen. Scale bar represents 20 µm (40x) and 200 µm (5x). **F-I)** Representative immunofluorescence images and quantitative analyses for **(F&H)** p21 (red) and **(G&I)** HMGB1 (green) in arteries from four groups of mice. L represents lumen. Scale bar, 20 µm. n = 5 independent biological samples. Data shown as mean ± SD.

Then, we evaluated whether this transcriptional shift was accompanied by reduced lipid accumulation in vivo, using Sudan Black B (SBB) staining **(Fig. 7C)**. While WT-Y mice displayed minimal staining (4.8 ± 3.3%), both aged C57BL/6 J (WT-O, 24 month old) and LAKI mice exhibited pronounced lipid deposition within the vascular wall, as evidenced by intense dark-blue staining (WT-O: 13.0 ± 1.4, p < 0.01; LAKI: 11.7 ± 5.4, p < 0.05) **(Fig. 7D)**. Notably, CHBA treatment significantly reduced the area of vascular lipid accumulation within the vascular wall in LAKI mice (LAKI+C: 2.8 ± 2.1%, p < 0.05) to a level comparable to that in WT-Y mice (p > 0.05). These findings suggest that CHBA alleviates age-associated lipid deposition in the vasculature.

To further test if treatment further alleviates senescence hallmarks, we assessed the level of SA-β-Gal on aortic cross-sections. The vascular wall of LAKI mice showed positive blue staining, which was markedly reduced after CHBA treatment at both low (5×) and high (40×) magnifications **(Fig. 7E)**. This result was further corroborated by immunostaining for additional senescence markers. Specifically, immunofluorescence staining revealed that p21 was markedly elevated in aged WT-O arteries, with significant abundance of p21+ cells in LAKI vessels (41.6 ± 15.6%); however, this effect was significantly reduced after CHBA treatment (16.9 ± 12.9% and p <0.05) **(Fig. 7F, H)**. We also evaluated high mobility group box 1 (HMGB1), a nuclear protein that is actively released extracellularly to promote SASP activation during aging and inflammation^57^ **(Fig. 7G)**. In young vessels, HMGB1 was predominantly localized within the nucleus as evidenced by a high percentage of positive HMGB1 nuclear area (44.8 ± 8.3%), while both WT-O (16.1 ± 3.7%, p < 0.0001) and LAKI groups (15.7 ± 4.1%, p < 0.0001) displayed significantly reduced nuclear HMGB1 signal **(Fig. 7I)**. Importantly, CHBA treatment significantly increased the nuclear accumulation of HMGB1 in the vascular wall as compared to LAKI controls (28.8 ± 7.5%, p < 0.05), restoring nuclear HMGB1 localization to levels closer to WT-Y arteries (p < 0.01). Overall, these results indicate that CHBA mitigates vascular aging in vivo, alleviating lipid accumulation and reducing senescent cell accumulation in the vascular wall.

### 8. Administration of CHBA modulates vascular structure and promotes ECM integrity

To investigate the functional relevance of the aging-associated genes that were elevated by CHBA treatment, we identified genes that were downregulated in the progeroid vasculature of LAKI mice relative to age-matched WT-Y controls and subsequently upregulated by CHBA treatment. To this end, we generated a ‘recovered-up’ gene set comprising the intersection of DEGs that were downregulated in LAKI vasculature (or equivalent to upregulated in WT-Y vs. LAKI, n=294) with DEGs upregulated in LAKI+C vs LAKI (n=396) **(Fig. 8A)**. GO analysis of these overlapping genes (n=156) revealed enrichment of pathways associated with ECM organization (‘cell-matrix adhesion’, ‘extracellular structure/ECM organization’, ‘basement membrane organization’ and ‘elastic fiber assembly’) and vascular remodeling processes (‘connective tissue development’, ‘regulation of blood circulation’, ‘vascular process in circulatory system’ and ‘artery development’) **(Fig. 8B)**.

**Figure 8:**
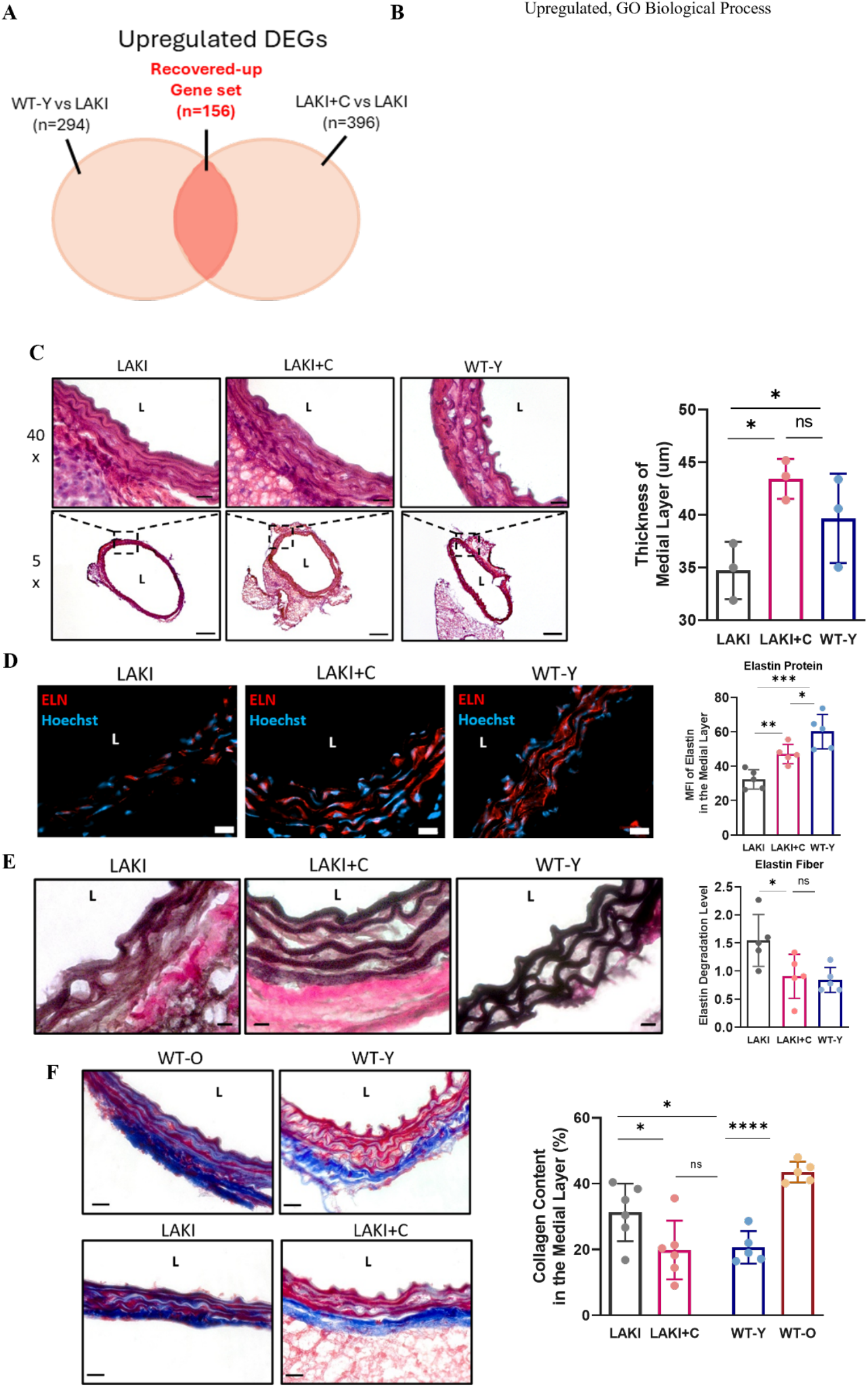
Administration of CHBA modulates vascular physiological structure and promotes ECM integrity in arteries of LAKI mice. **A)** Venn diagram of the recovered-up gene sets. **B)** GO enrichment analysis of recovered-up gene sets, highlighting Biological Processes associated with ECM and vascular development. Bubble size represents gene count. **C)** Quantification of the thickness of medial layer based on the hematoxylin and eosin staining, n=3 biological replicates. Data shown as mean ± SD. **D)** Immunostaining and quantitative analysis of elastin (ELN) in arteries of three groups of mice: LAKI, LAKI+C and WT-Y. L represents lumen. Scale bar, 20 μm. n = 5 independent biologically samples. Data shown as mean ± SD. **E)** Representative images of Verhoeff staining in cross sections from arteries of three groups. Scale bar represents 20 µm. L represents lumen. Quantification of the elastin degradation score. n=5 independent biological samples. Data shown as mean ± SD. **F)** Representative images of Trichrome staining in cross sections from arteries of four groups of mice: WT-O, WT-Y, LAKI and LAKI+C and quantification of collagen content in the medial layer. Scale bar represents 20 µm. L represents lumen. n=5 independent biological samples. Data shown as mean ± SD.

We next examined whether these transcriptional changes were accompanied by improved preservation of vascular wall structure during aging. Medial wall thickness was assessed in cross-sections of aortic tissue using hematoxylin and eosin (H&E) staining **(Fig. 8C)**. As expected, LAKI mice exhibited significantly thinner vessel walls (34.7 ± 2.7 µm) compared to WT-Y mice (39.6 ± 4.3 µm, p < 0.05). In contrast, LAKI+C mice displayed a marked increase in medial wall thickness compared to untreated LAKI mice (43.4 ± 1.9 µm, p < 0.05), reaching values closer to those of young mice (p > 0.05).

We further examined whether CHBA could preserve key ECM components, particularly elastin and collagen, which undergoes significant changes during aging. Immunofluorescence staining for elastin (ELN) revealed a marked increase in the ELN^+^ mean fluorescence intensity (MFI) in the medial layer of LAKI+C vessels compared to untreated LAKI arteries **(Fig. 8D)**. Quantitative analysis confirmed 1.4-fold increase in elastin protein expression following CHBA treatment relative to untreated control (p < 0.01), bringing the level of ELN expression closer to WT-Y mice (WT-Y: 1.8-fold higher compared to LAKI, p < 0.001; also 1.2-fold higher compared to LAKI+C, p < 0.05).

The integrity of elastin fibers was further evaluated using Verhoeff staining^58^. In LAKI mice, elastin fibers appeared fragmented and disorganized, whereas vessels from CHBA-treated LAKI mice (LAKI+C) exhibited continuous, well-aligned elastin fibers **(Fig. 8E)**. Elastin fiber integrity was quantified using a semiquantitative scoring system (0 = intact, 3 = severely degraded). As shown in **Supplementary Fig. S5**, CHBA treatment significantly attenuated elastin fiber degradation (LAKI: 1.5 ± 0.4; LAKI+C: 0.9 ± 0.3, p < 0.05), restoring elastin fiber integrity to levels comparable to those of young WT mice (WT-Y: 0.8 ± 0.2, p > 0.05 compared to LAKI+C).

Since elastin degradation is often accompanied by collagen deposition during vascular aging^59^, we assessed collagen content using Masson’s trichrome staining, visualized as blue-stained regions **(Fig. 8F)**. In aged WT-O and LAKI arteries, collagen deposition was evident throughout the vascular wall. In contrast, CHBA-treated progeroid mice displayed reduced collagen deposition and well-preserved, red-stained muscle layers, resembling the vascular architecture of young WT mice. Quantitative analysis confirmed that CHBA significantly reduced medial collagen content from 31.3 ± 8.4 % in LAKI mice to 19.8 ± 8.8 % in LAKI+C mice (n = 5, p < 0.05), restoring collagen levels to those observed in young WT vessels (WT-Y: 20.6 ± 4.9 %, p > 0.05, compared to LAKI+C). Taking together, these results suggest that CHBA treatment preserves vascular ECM integrity in progeroid mice by reducing pathological collagen accumulation and maintaining elastin fiber architecture.

### 9. Administration of CHBA promotes vascular remodeling in progeroid mice

Similarly, GO analysis of the aging-associated genes that were upregulated by CHBA (**Fig. 8A**, n=156) revealed enrichment of pathways related to actin filament organization (‘actin filament/ contractile actin filament bundle assembly’ and ‘actin filament organization’) and VSMC functions (‘smooth muscle contraction’, ‘negative regulation o smooth muscle cell migration’, ‘regulation of smooth muscle cell proliferation’ and ‘smooth muscle tissue development’) **(Fig. 9A)**. To further define the effect of CHBA treatment at the cell-type level, we performed Gene Set Enrichment Analysis (GSEA) on the VSMC cluster comparing LAKI+C and LAKI mice. REACTOME pathway analysis revealed significant downregulation of pathways associated with extracellular matrix remodeling, including “ECM proteoglycans”, “extracellular matrix organization”, “cell membrane–ECM interactions”, and “collagen biosynthesis and crosslinking” **(Fig. 9B)**. In parallel, HALLMARK pathway analysis identified downregulation of pathways are associated with well-established proatherogenic functions: “Interferon gamma response” (NES = -1.57, p.adj< 0.01) and “Adipogenesis” (NES = -1.82, p.adj< 0.0001) **(Fig. 9C)**. Meanwhile, pathways upregulated in LAKI+C VSMCs included mRNA splicing, epigenetic regulation of gene expression, cell junction organization and cell-cell communication, suggesting transcriptional remodeling with improved cell-cell junction and communication following CHBA treatment **(Fig. 9B)**. To further examine changes in intercellular communication, we applied CellChat analysis to the snRNA-seq dataset. Compared to LAKI, LAKI+C mice showed increased number of intercellular interactions **(Fig. 9D)**, although they remained slightly lower than those observed in WT-Y arteries **(Supplementary Fig. S6)**. The increase in intercellular communication involved multiple vascular cell populations, including VSMCs, ECs, fibroblasts, adipocytes, and neural cells. Among these, VSMCs emerged as the predominant signaling (sender) population following CHBA treatment, while ECs also displayed increased signaling activity.

**Figure 9:**
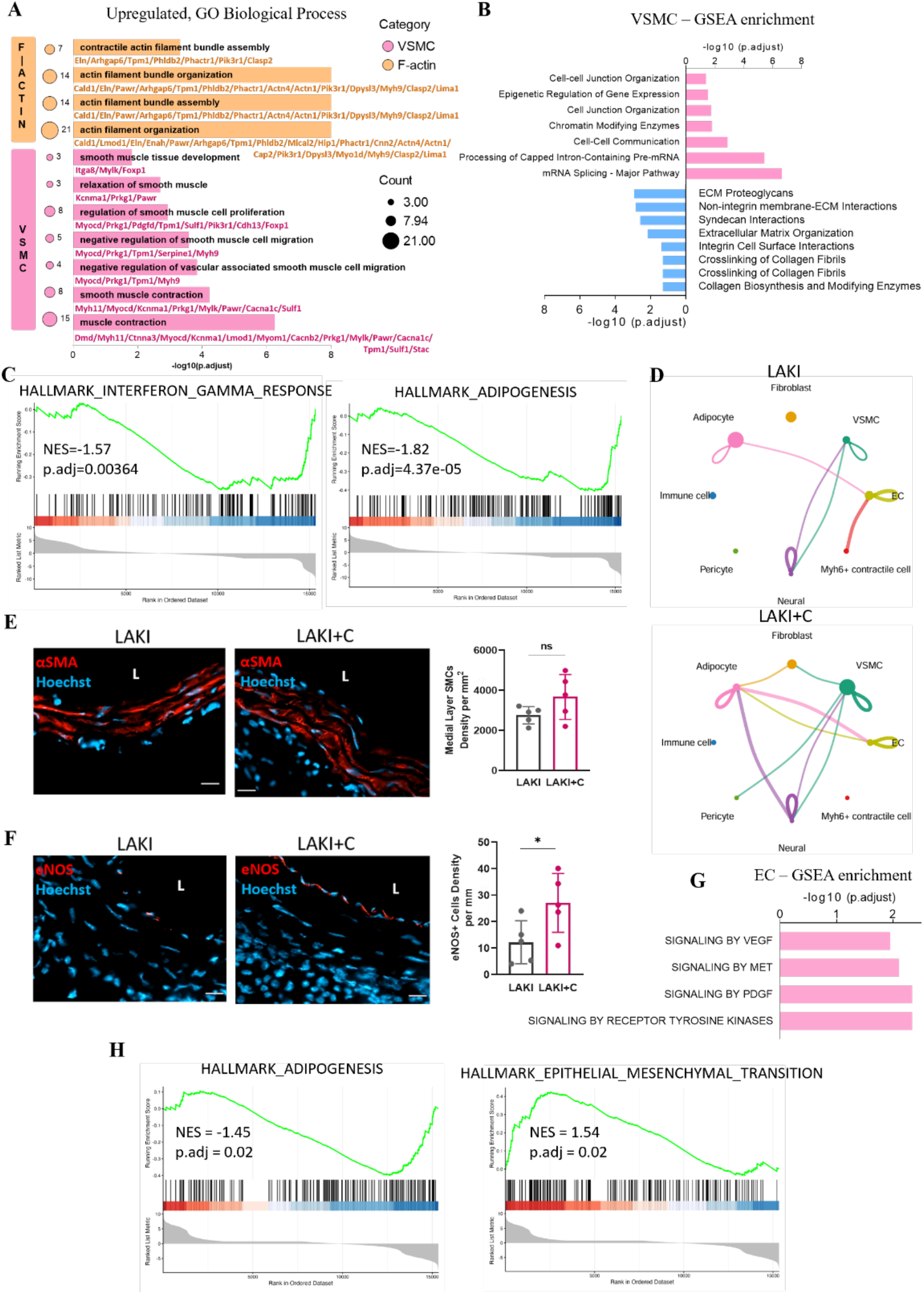
Administration of CHBA modulates vascular physiological structure in arteries of progeroid mice. **A)** GO enrichment analysis of rescued-up gene sets, highlighting Biological Processes associated with actin filament (F-actin) and VSMC. Bubble size represents gene count. **B)** Significantly enriched REACTOME pathway from GSEA in VSMCs cluster from LAKI+C mice compared to LAKI. Upregulated pathways are shown in pink (P adj. <0.05), downregulated pathways are shown in blue (P adj. <0.05). **C)** The visualization of significantly enriched hallmark terms from GSEA (p.adj < 0.05) in VSMCs from arteries following CHBA treatment. **D)** CellChat analysis showed the number of interactions between pairwise cell populations among all cell clusters in LAKI (top) and LAKI+C (bottom) groups. Each node represents a cell type, with node size proportional to the total number of interactions associated with that population. Connecting lines represent predicted ligand–receptor-mediated communication between cell types. Line color corresponds to the sender cell population and line thickness indicates the relative strength of communication. **E)** Immunostaining and quantitative analysis of αSMA (red) in arteries from LAKI and LAKI+C mice. L represents lumen. Scale bar, 20 µm. n = 5 biologically independent samples. Data shown as mean ± SD. **F)** Immunostaining and quantitative analysis of eNOS (red) in arteries from LAKI and LAKI+C mice. L represents lumen. Scale bar, 20 µm. n = 5 independent biological samples. Data shown as mean ± SD. **G)** Significantly enriched REACTOME pathway from GSEA in ECs cluster from LAKI+C mice compared to LAKI. Upregulated pathways are shown in pink (P adj. <0.05). **H)** Visualization of significantly enriched hallmark gene sets (p.adj < 0.05) in ECs from arteries following CHBA treatment.

We further investigated whether these transcriptomic changes affect the vascular integrity in vivo, by assessing the density of EC and SMC using endothelial nitric oxide synthase (eNOS) and αSMA immunostaining, respectively. The density of αSMA⁺ cells in the medial layer showed a modest increase with CHBA treatment but did not reach statistical significance (LAKI: 2757.3 ± 432.4 αSMA⁺ cells per mm^2^; LAKI+C: 3671.5 ± 1116.8 αSMA⁺ cells per mm^2^, p > 0.05) **(Fig. 9E)**. Notably, compared to untreated LAKI mice, CHBA treatment significantly increased the density of eNOS+ cells along the luminal surface of arteries (LAKI: 12.1 ± 8.0 eNOS+ cells per mm; LAKI+C: 27.0 ± 11.1 eNOS+ cells per mm, p < 0.05), indicating enhanced endothelial integrity **(Fig. 9F)**. To further examine endothelial changes at the transcriptional level, we performed GSEA analysis in the EC cluster. REACTOME pathway analysis comparing LAKI+C and LAKI mice identified a broad set of upregulated endothelial signaling pathways, including Signaling by VEGF, MET, Platelet-derived growth factor (PDGF), and receptor tyrosine kinases **(Fig. 9G)**. Consistent with the findings in the VSMC cluster, HALLMARK pathway analysis of ECs revealed significant suppression of the adipogenesis pathway (NES = - 1.45, p.adj<0.05), along with enrichment of epithelial to mesenchymal transition pathway (NES = 1.54, p.adj< 0.05), which may be associated with tissue regeneration and repair **(Fig. 9H)**. Collectively, these findings suggest that CHBA partially restores vascular cell function in progeroid arteries by promoting contractile features in progeroid arteries, attenuating ECM remodeling, enhancing intercellular communication, and preserving endothelial integrity.

## Discussion

Our study identified GPR81 signaling as a regulator of vascular senescence and vascular homeostasis. We found GPR81 to be essential for maintaining lipid and energy homeostasis as well as vascular cell senescence, as evidenced by lipid accumulation, ferroptotic stress, mitochondrial dysfunction and senescent-associated hallmarks in cultured VSMCs and ECs upon GPR81 knockdown. Importantly, EC-specific deletion of GPR81 in young mice was sufficient to induce age-associated changes in aortic tissue. Conversely, pharmacological activation of GPR81 restored mitochondrial and lipid metabolism as well as alleviated ferroptotic stress and cellular senescence in vitro and in vivo. In addition, GPR81 activation preserved endothelial integrity and improved ECM organization in progeroid mice. Together, these findings indicate that GPR81 signaling may be a promising therapeutic target for mitigating vascular aging and related cardiovascular disease.

Our results demonstrated that vascular senescence is associated with lipid dysregulation and enhanced ferroptotic stress. Specifically, senescent vascular cells exhibited increased CD36 expression, accompanied by enhanced lipid accumulation and lipid peroxidation. These findings are consistent with previous studies linking cellular senescence to lipid accumulation and impaired endothelial function^60–63^. Moreover, age-related cardiovascular diseases, including atherosclerosis and other arterial pathologies, have been attributed in part to the progressive accumulation of lipids within the vascular wall^64^. CD36 has also been extensively implicated in mediating excessive lipid uptake from the extracellular environment while promoting pro-inflammatory signaling.^65–67^ In parallel, we observed activation of pro-ferroptotic signaling, a process closely associated with lipid peroxidation and previously shown to occur in senescent vascular smooth muscle cells (VSMCs) and aged arteries^20,23,26^. Consistent with these reports, senescent vascular cells in our study exhibited increased ferroptotic stress accompanied by pronounced lipid peroxidation, suggesting that age-related lipid dysregulation may sensitize vascular cells to ferroptotic stress.

GPR81 is a well-known key regulator of lipid metabolism in adipose tissue, where it suppresses lipolysis via inhibition of the cAMP–PKA axis^28,29^. Previous studies demonstrated that pharmacological activation of GPR81 influences systemic lipid metabolism, as evidenced by suppression of fasting plasma FFA levels and improved insulin sensitivity in mice^32^, suggesting that GPR81 may exert boarder metabolic effects beyond adipocytes. Indeed, GPR81 was found to be expressed in the cardiovascular system^31^, and cAMP-PKA signaling has been implicated in the regulation of lipolysis in ECs^68^, yet the direct role of GRP81 in vascular cells remains largely unexplored. Our findings demonstrated that GPR81 signaling contributes to lipid metabolism in vascular cells, as GPR81 knockdown led to intracellular neutral lipid accumulation, enhanced lipid uptake and impaired β-oxidation. Conversely, pharmacological activation of GPR81 by the selective agonist CHBA reduced lipid accumulation, and partially restored FA oxidation in senescent vascular cells. Notably, FABP3, a clinically relevant marker for coronary and peripheral artery disease^41^, was among the most responsive to CHBA. In addition, FATP1, FATP4, and FATP6, which facilitate the uptake of long-chain fatty acids (LCFAs) and possess acyl-coenzyme A (CoA) synthetase (ACS) activity that promotes intracellular polyunsaturated fatty acid (PUFA) metabolism^10^, were all upregulated by CHBA treatment. On the other hand, the role of GPR81 in ferroptosis remains unclear and perhaps context dependent, since GPR81 inhibition amplified the susceptibility of cancer cells to ferroptosis by downregulating SCD1^69^, but suppressed ferroptosis in alveolar epithelial cells^70^. Similar to cancer cells, we found that activation of GPR81 signaling alleviated ferroptotic stress in senescent ECs and VSMCs. Collectively, our data implicate GPR81 signaling in the regulation of vascular lipid metabolism and the suppression of lipid peroxidation-driven ferroptotic stress.

Our results also show that GPR81 knockdown induced vascular senescence, as evidenced by elevated SA-β-gal activity, reduced proliferation (Ki67), increased DNA damage (γH2AX) and impaired mitochondrial metabolism in vitro. Notably, these findings are supported by an EC-specific GPR81 knockout mouse model, which provides compelling evidence that GPR81 plays an essential role in maintaining vascular homeostasis and preventing vascular senescence. In young (19-week-old) mice, EC-specific deletion of GPR81 increased senescence markers throughout the aortic wall, likely as a consequence of impaired endothelial barrier function, reduced nitric oxide signaling, and the production of pro-inflammatory paracrine factors. Moreover, Hcar1^fl/fl^ Cdh5-Cre mice exhibited age-like pathological changes in the aorta, including disrupted endothelial junctional integrity and elastin disorganization. These findings support the concept that endothelial dysfunction is sufficient to initiate pathological remodeling and senescence throughout the vascular wall. Interestingly, GPR81 protein decreased significantly in the aortas of both naturally aged and progeroid mice, whereas GPR81 signaling was reduced in the gastrocnemius (GA) without a comparable decline in protein expression^71^. These findings suggest that age-associated alterations in GPR81 signaling vary across tissues, with a particularly pronounced effect in the vasculature. Collectively, these observations underscore the importance of vascular GPR81 signaling and identify it as a promising therapeutic target for mitigating age-related vascular dysfunction.

Furthermore, pharmacological activation of GPR81 alleviated the hallmarks of cellular senescence and reprogrammed cellular metabolism. Consistent with this, some studies have shown that exercise-induced lactate signaling through GPR81 promotes angiogenesis and improves arterial function^72,73^, indicating a broader role for GPR81 in vascular homeostasis. Although the underlying mechanism remains unclear, we found that GPR81 activation induced metabolic reprogramming by reducing oxidative phosphorylation and enhancing glycolytic flux in both senescent vascular cell types, as evidenced by decreased oxygen consumption together with increased glycolysis and glycolytic capacity. ECs rely primarily on glycolysis for ATP production, and disruption of the key glycolytic enzyme, PFKFB3 impairs endothelial migration and vessel formation^74,75^. Indeed, CHBA enhanced EC migration, which was impaired by GPR81 knockdown. In contrast, VSMCs in their contractile state are more dependent on mitochondrial oxidative metabolism, while phenotypic switching from the contractile to the synthetic phenotype - migratory and proliferative - is associated with increased reliance on glycolysis^76^, suggesting that GPR81 activation might have promoted VSMC switch towards the synthetic phenotype. In this context, the reduction in oxidative phosphorylation may represent an adaptive response that limits mitochondrial ROS production to combat oxidative stress^77,78^. Overall, these findings indicate that metabolic reprogramming of vascular cells mediated by GPR81 may contribute to the attenuation of vascular cell senescence.

Consistent with our in vitro observations, administration of CHBA to progeroid mice suppressed lipid-associated pathways, alleviated lipid deposition and reduced accumulation of senescent vascular cells in the mouse aorta. Indeed, snRNA-seq analysis revealed that CHBA partially shifted the arterial transcriptomic profile of progeroid mice toward that of young (3-month-old) WT mice. Notably, CHBA also remodeled the cellular composition of the progeroid vascular wall by reducing the proportion of adipocytes while restoring the proportion of vascular smooth muscle cells (VSMCs) to levels comparable to those of young arteries. These changes were accompanied by enhanced intercellular communication, increased endothelial signaling, and elevated eNOS expression in luminal endothelial cells (ECs). This finding is particularly important because eNOS is the primary source of vascular nitric oxide (NO), which is essential for maintaining vasodilation, vascular redox homeostasis, and endothelial function while protecting against vascular aging^79^. These observations are also consistent with previous reports demonstrating that GPR81 signaling protects ECs from oscillatory shear stress (OSS)-induced inflammation^33^.

In addition, snRNA-seq analysis revealed that CHBA treatment enriched pathways involved in vascular remodeling and ECM organization, processes that are essential for maintaining vascular homeostasis and preventing age-associated vascular diseases such as atherosclerosis^80,81^. Indeed, CHBA-treated progeroid arteries showed decreased collagen deposition and reduced elastin fragmentation, effectively maintaining healthy ECM balance and structural integrity of the vascular wall. Excessive collagen accumulation and elastin degradation are hallmarks of vascular aging that promote arterial stiffening, impair vascular compliance, and contribute to vascular dysfunction^58,59,82^.

Ultimately, CHBA treatment reduced the expression of the cell cycle arrest marker p21 and SA-β-gal activity while promoting the nuclear retention of HMGB1, which normally translocate from the nucleus to the cytoplasm during cellular senescence and inflammation^57^. Interestingly, we did not detect intracellular cytoplasmic accumulation of HMGB1 in untreated progeroid arteries, consistent with previous reports showing that cytoplasmic HMGB1 is rapidly secreted into the extracellular space rather than retained intracellularly^83^. Collectively, these findings identify GPR81 signaling as a critical regulator of vascular remodeling during aging and suggest that targeting GPR81 may represent a promising therapeutic strategy for mitigating ECM-driven vascular stiffening and the progression of age-associated vascular disease.

Finally, several limitations of this study should be acknowledged. First, although the Lmna^G609G^ knock-in mouse model recapitulates many features of Hutchinson–Gilford progeria syndrome, including severe cardiovascular complications^84^, it does not fully reflect the physiological aging process. Therefore, additional studies in naturally aged mice are warranted to determine whether GPR81 activation confers similar vascular benefits during normal aging. Furthermore, CHBA was administered systemically via intraperitoneal injection. Because GPR81 is expressed in multiple tissues and cell types, CHBA may exert effects on organs beyond the vasculature, such as the liver, which could indirectly influence vascular responses. Future studies using vascular cell-specific genetic models or targeted drug delivery approaches will be important to define the cell-autonomous role of GPR81 signaling in senescent vascular cells.

## Materials and Methods

### 1. Primary human vascular cell culture

Primary HASMCs and HUVECs obtained from American Type Culture Collection (ATCC). VSMCs were cultured in DMEM (Gibco) containing 10% (v/v) fetal bovine serum (FBS) (Atlanta Biologicals) and 1% (v/v) Antibiotic-Antimycotic cocktail (Thermo Fisher Scientific). HUVECs were cultured in Endothelial Cell Growth Medium-2 (EGM-2; CC-3162, Lonza). The cells were incubated at 37 °C with 5% CO_2_ in a humidified atmosphere. All cells were routinely tested for mycoplasma contamination, that was prevented by employing 5 µg/ml Plasmocin® prophylactic (ant-mpp, InvivoGen) in culture media. For studying replicative senescence in vitro, cells at passages 3-6 were used as EPs control, while passages 10-14 were used as replicative senescence cells, LPs.

### 2. shRNA vectors for *GPR81* knockdown

MISSION® shRNA bacterial glycerol stock that carried either the empty vector (pLKO.1-Puro, SHC001, Sigma-Aldrich) or GPR81-specific shRNA (NM_032554, TRCN0000008942, Sigma-Aldrich) were purchased. Bacteria glycerol stocks were expanded, and plasmids were extracted using the NucleoBond Xtra Midi kit (Cat # 740410, Macherey-Nagel).

### 3. Lentivirus production and transduction

Lentivirus was produced in 293T cells with three plasmids (shRNA encoding lentiviral vector, psPAX2, and pMD2.G) using the standard calcium phosphate precipitation method^85^. EPs VSMCs and HUVECs were transduced with lentiviral particles that encoded the puromycin resistance gene and carried either the empty vector (Y_Empty) or shRNA targeting GPR81 (Y_shGPR81). Cells were selected with 1 µg/mL puromycin (Cat # BML-GR312, Enzo Life Sciences) for 5 days before used for future assays.

### 4. RNA isolation, cDNA synthesis and Quantitative real-time PCR

Total RNA was extracted from cultured cells using RNeasy Plus Mini Kit (Cat # 74134, Qiagen) according to the manufacturer’s instruction. One microgram of isolated RNA per sample was reverse transcribed to cDNA with the High-Capacity cDNA Reverse Transcription Kit (Cat # 4368814, Applied Biosystems). To assess gene expression, real-time PCR was performed using Bio-Rad CFX96 system (Bio-Rad Laboratories) with SYBR Select Master Mix (Cat # A25742, Applied Biosystems). The primer sequences were listed in **Supplementary Table 1**. Cycle threshold (CT) values were recorded, and then relative gene expression levels were calculated by the ΔΔC_t_ method to quantify fold changes of the interested genes and internally normalized to the cycle number of RPL32.

### 5. Protein extraction and immunoblotting

For protein extraction, cells were lysed in RIPA lysis buffer (Cat#89900, Thermo Fisher Scientific) supplemented with 1X Halt Protease Inhibitor and 41.67 mM dithiothreitol (Cat # 14265, DTT; Cell Signaling Technology). Besides, artery tissues were lysed in the same buffer by bead disruption in bead lysis tubes (Cat # GREENR5-RNA, Next Advance) using the Bullet Blender® Gold tissue homogenizer (Next Advance). Protein concentration of samples was determined using the BCA protein assay kit (Cat# 23250, Thermo Fisher Scientific). For immunoblotting, lysates were centrifuged and 1X blue loading dye (3X; Cat# 7722S, Cell Signaling Technology) was added. Protein was denatured by incubation at 95 °C for 5 min, separated by SDS-PAGE using 4-20% Tris-Glycine gels (Cat# XP04205BOX, Thermo Fisher Scientific) at 120 V for 1.5 hour at room temperature (RT), transferred to nitrocellulose membranes (Cat# 1620113, Bio-Rad Laboratories) and blocked in 5% (w/v) nonfat dry milk in Tris-buffered saline (20 mM Tris, 150 mM NaCl) with 0.1% (v/v) Tween® 20 detergent (TBST) for 1 hour at RT. Membranes were blotted with primary antibody (anti-GPR81, # PA5-67873, Invitrogen, 1:1000 in blocking buffer) overnight at 4°C followed by incubation with horseradish peroxidase–conjugated anti rabbit IgG secondary antibody (Cell Signaling Technology, 1:1000) for 1 hour at RT.

Immunoreactive proteins were visualized using SuperSignal West Pico PLUS chemiluminescence substrate (Cat # 34578, Thermo Fisher Scientific). Protein bands were captured using the ChemiDoc MP imaging system (Bio-Rad), and the relative expression level of proteins were quantified using the Bio-Rad Image Lab 6 software.

### 6. Senescence-associated β-galactosidase assay (SA-β-Gal)

Cellular accumulation of β-galactosidase activity at pH=6 is a well-established marker of senescence. The senescent status of HUVECs and VSMCs were detected by a senescence detection kit (Cat# 9860, Cell Signaling) according to the manufacturer’s instruction. Senescent cells in culture were assessed after 24 hour incubation with freshly prepared staining solution at 37 °C, while senescent cells accumulated in artery tissue sections were detected after 48 hour incubation. The rate of positively stained cells (blue cells) to total cells was calculated in 5 randomly chosen microscopic fields using the Zeiss Axio Observer Z1 (Oberkochen, Germany) inverted microscope with an ORCA-ER CCD camera (Hamamatsu, Japan). The level of senescent vascular cells in the vascular wall was determined by the percentage of positive stained areas in the wall.

### 7. Detection of ROS in live cells

To visualize intracellular reactive oxygen species (ROS) in human vascular cells in culture, ROS content was measured by DCFDA/H2DCFDA–Cellular ROS Assay Kit (Cat # ab113851, Abcam) according to the manufacturer’s instructions. Cells were treated with DCFDA (20 µM) for 20 min at 37 °C in live imaging solution (LIS) and then stained with the Hoechst 33342 nuclear dye at 1:500 dilution in LIS for 5 min at 37°C. Followed by washing cells thrice with LIS, images were taken using the Zeiss Axio Observer Z1 imaging system.

### 8. Labile iron pool assay

After washing thrice with PBS, cultured cells were incubated with 1 µM FerroOrange probe (Dojindo, Japan) for 30 min at 37 °C. The fluorescence intensity of Alexa Fluor 568 was monitored using the Zeiss Axio Observer Z1 imaging system. Images were quantified for approximately 100 cells/ sample from 5 randomly chosen microscopic fields. The FerroOrange fluorescent intensity is proportional to the labile iron concentration.

### 9. Visualization of cellular lipid contents using BODIPY

Cultured cells were washed thrice with PBS, then incubated in 5 µM BODIPY dye in LIS for 25 min at 37°C. Cells were then stained with the Hoechst 33342 nuclear dye at 1:500 dilution in LIS for 5 min at 37°C. This was followed by washing cells thrice with LIS and monitored under the Zeiss Axio Observer Z1 imaging system. BODIPY 493/503 (Cat # D3922, Thermo Fisher Scientific), which specifically bind to neutral lipids, was used to visualize neutral lipid droplets accumulation in the cytoplasm. BODIPY 581/591 C11 lipid peroxidation sensor (Cat # D3861, Thermo Fisher Scientific) was used to detect lipid peroxides.

### 10. Immunocytochemistry and immunohistochemistry

For immunocytochemistry, cells were washed with cold PBS (4°C) once and fixed for 10 min with 4% (w/v) paraformaldehyde, followed by permeabilization with 0.1% (v/v) Triton X-100 in PBS for 10 min at RT. Next, samples were blocked using blocking buffer [5% (v/v) goat serum (Life Technologies) in PBS with 0.01% (v/v) Triton X-100] for 1 hour at RT and then incubated with primary antibodies **(Supplementary Table 2)** diluted in blocking buffer at 4°C overnight. Subsequently, cells were washed three times and incubated with Alexa Fluor 488- or 568-conjugated anti-mouse/rabbit secondary antibodies (1:200 in blocking buffer, Invitrogen) for 1 hour at RT. Nuclei were stained using Hoechst 33342 for 5 min at RT (Thermo Fisher Scientific).

Immunohistochemistry was carried out as follows: Tissue sections were washed three times in PBS to remove the optimal cutting temperature (OCT) embedding medium (Sakura Finetek). Subsequently, they were treated with pre-warmed permeabilization buffer [0.1 % (v/v) triton X-100 in 1X PBS] for 10 min. Next, tissue sections were blocked with blocking buffer at RT for 1 hour followed by incubation at 4 °C overnight with primary antibodies. Following three washes with wash buffer [0.01 % (v/v) triton X-100 in 1X PBS], tissue sections were treated with 0.3 % hydrogen peroxide (VWR Chemicals) for 10 min to suppress endogenous peroxidase activity. Next, they were washed with wash buffer again twice and incubated for 1 hour with Alexa Fluro 488- or 568-conjugated anti-mouse/rabbit secondary antibodies (1:200, Invitrogen). The sections were mounted with ProLong™ Glass Antifade Mountant with NucBlue™ Stain (Thermofisher). Image acquisition was performed by using the Zeiss Axio Observer Z1 imaging system. Primary antibodies were listed in **Supplementary Table 2.**

### 11. Cellular bioenergetics - measurement of mitochondria respiration and glycolysis

The oxygen consumption rate (OCR), as an indicator of mitochondrial respiration, and the extracellular acidification rate (ECAR), as a measure of glycolytic activity, were quantified using the Seahorse XFe96 Analyzer (Agilent). HUVECs were seeded in Seahorse XF96 cell culture microplates (Cat# 103794-100, Agilent) at a density of 1.2 × 10⁴ cells per well, and VSMCs were seeded at 8 × 10³ cells per well for overnight. Prior to the assay, the culture medium was gently removed and replaced with Seahorse XF DMEM (Agilent) supplemented as follows: for OCR measurements, 10 mM glucose, 1 mM pyruvate, and 2 mM glutamine; for ECAR measurements, 2 mM glutamine. Cells were then equilibrated for 1 hour at 37 °C in a non-CO₂ incubator. For OCR analysis, oligomycin (1.5 μM), FCCP (1 μM), and a rotenone/antimycin A mixture (0.5 μM each) were injected sequentially. For ECAR analysis, glucose (10 mM), oligomycin (1 μM), and 2-deoxy-D-glucose (2-DG; 50 mM) were sequentially injected. All measurements were acquired and analyzed using the Seahorse Wave software.

### 12. Cellular bioenergetics – measurement of FAO

The ability of cells to utilize endogenous fatty acids was assessed by monitoring changes in OCR. HUVECs were seeded in Seahorse XF96 cell culture microplates at a density of 1 × 10⁴ cells per well, and VSMCs were seeded at 6 × 10³ cells per well and allowed to adhere overnight. The following day, culture medium was removed and replaced with substrate-limited medium [Seahorse XF DMEM (Cat# 103575-100, Agilent) plus 1% FBS, 0.5 mM glucose (Cat# 103577-100, Agilent), 0.5 mM L-carnitine (Cat# 0158, Sigma-Aldrich), and 1 mM glutamine (Cat# 103579-100, Agilent)], and cells were incubated overnight. On the day of the assay, the substrate-limited medium was replaced with oxidation medium [Seahorse XF DMEM (Cat# 103575-100, Agilent) supplemented with 2 mM glucose (Cat# 103577-100, Agilent) and 0.5 mM L-carnitine (Cat# 0158, Sigma-Aldrich)], and cells were equilibrated for 45 min at 37 °C in a non-CO₂ incubator. Cells were then incubated for 15 min in oxidation medium containing either etomoxir (4 μM) or vehicle control. OCR was subsequently measured following sequential injections of oligomycin (1.5 μM), FCCP (1 μM), and a rotenone/antimycin A mixture (0.5 μM each), according to the XF Cell Mito Stress Test protocol. Data acquisition and analysis were performed using Seahorse Wave software.

### 13. Cell scratch assay

To investigate the migration ability of HUVECs, cells were seeded in 24-well plates at a density of 1×10^5^ cells per well and allowed to reach confluence to form a uniform monolayer. A linear scratch was generated across the cell layer using a sterile pipette tip. Immediately after scratching, images were captured to document the initial wound area (0 hour) by Zeiss Axio Observer Z1 inverted microscope with an ORCA-ER CCD camera. Cells were then cultured and wound closure was imaged again after 6 and 24 hours. The migration ability was quantified using ImageJ software. Migration ability was calculated as: [(A_0_ – A_t_)/A_0_] × 100%, where A_0_ represents the initial wound area at 0 hour, and A_t_ represents the remaining wound area at each subsequent time point.

### 14. Administration of 3-chloro-5-hydroxy BA (CHBA) in mice and tissue collection

Lamin-A Knocked In (LAKI, C57BL/6-LmnaG609G/G609G) mice were intraperitoneally injected with CHBA [10 mg/kg dissolved in 5% (v/v) DMSO in corn oil (Cat # S6701, Selleckchem Houston); abbreviated as LAKI+C: 7 males and 6 females) or vehicle control [5% (v/v) DMSO in corn oil; LAKI: 7 females and 6 males), three times per week for 4 weeks. Young WT mice (WT-Y, C57BL/6J: 5 females, 5 males) were also administered vehicle [5% (v/v) DMSO in corn oil) and were used as controls. Following 4 weeks of treatment, mice were euthanized by carbon dioxide overdose followed by cervical dislocation. All research involving animals followed approved protocols from the Institutional Animal Care and Use Committee (IACUC) of the University at Buffalo. Aortas were harvested, perfused with cold PBS to remove residual blood, and dissected to remove surrounding adipose tissue. For immunohistochemistry analyses, aorta tissues were embedded in OCT compound by freezing in a dry ice/2-methylbutane bath. Additional aorta tissues were snap frozen in liquid nitrogen and stored at −80 °C until further analysis. Blood was collected at the time of euthanasia and centrifuged at 3,500 rpm for 15 min at 4 °C. The serum was carefully collected from the supernatant.

### 15. Endothelial cell specific Hcar1 deficient mice model generation

Hcar1^fl/fl^ mice were obtained from GemPharmatech (T058433), and Cdh5-Cre mice were obtained from The Jackson Laboratory (033055). All mice were on a C57BL/6J genetic background. Male and female mice at 19 weeks of age were used for experiments. To generate endothelial cell specific Hcar1 deficient mice, Hcar1^fl/fl^ mice were crossed with Cdh5-Cre mice as previously described^86^. Hcar1^fl/fl^ littermates were used as controls, and Hcar1^fl/fl^ Cdh5-Cre mice were used as endothelium Hcar1 deficient mice. All animal procedures were approved by and performed in accordance with the Institutional Animal Care and Use Committee of Fudan University (IDM2024016).

### 16. Histology (H&E, Trichrome, Verhoeff, and SBB staining)

Hematoxylin and Eosin (H&E) (Cat # ab245880, Abcam), Masson’s trichrome (Cat # ab150686, Abcam), Elastic-van-Gieson (EVG) (Cat # ab150667, Abcam) and Sudan Black B (SBB) (7 mg/ml in ethylene glycol, Sigma-Aldrich) were used to assess tissue morphology, collagen deposition, elastin degradation and lipid deposition, respectively. The aorta sections (10 μm) were stained according to the company’s protocols.

Elastic degradations were evaluated by an established elastin degradation score, which was adopted as follows: score 1, no elastin degradation, well-organized elastin lamina; score 2, mild elastin degradation with some interruptions or breaks in the lamina; score 3, moderate elastin degradation with multiple interruptions or breaks in the lamina; and score 4, severe elastin fragmentation or loss or aortic rupture.

### 17. Sample preparation for snRNA seq

Snap-frozen aortic tissues were thawed on ice for 5 min before processing. For each animal, 3 mg of aortic tissue was minced and lysed in digestion buffer [1 mM MgCl_2_, 5 M NaCl, 1 M Tris HCl and 0.1% (v/v) IGEPAL (Cat# I8896, Sigma-Aldrich) in PBS] by bead disruption in bead lysis tubes using the Bullet Blender® Gold tissue homogenizer. The lysate was then washed with wash buffer [2% (w/v) BSA in PBS] to remove the digestion buffer and preserve the nuclei integrity. The isolated nuclear suspension was passed through a cell strainer to remove debris and clumps, followed by centrifugation at 500 x g for 5 min at 4 °C. The nuclear pellet was resuspended in 1 ml wash buffer and centrifuged again at 500 x g for 10 min at 4 °C. After carefully removing the supernatant, the final nuclear pellet was resuspended in Minute^TM^ anti-clumping nuclei storage buffer (Invent Biotechnologies Inc.) with 200 U/ml RNase inhibitor (Cat# N8080119, Applied Biosystems) and processed for single-nucleus RNA sequencing.

### 18. snRNA-seq data analysis

The isolated nucleus from mice aorta were processed for snRNA sequencing at the University at Buffalo Genomics Core (UBGBC). The data were then processed using the standard Seurat (v5.0.0) workflow. Nuclei were filtered for mitochondria transcripts < 10%, nCount_RNA > 200 and nFeature_RNA > 25 to remove low quality nuclei, potential doublets and damaged nuclei. Data were normalized using SCTransform, followed by principal component analysis using RunPCA. To minimize sample variation, datasets were integrated using Harmony integration method. Clustering was then performed using FindNeighbors and FindClusters Seurat commands at a resolution of 0.5. Uniform manifold approximation and projection (UMAP) was performed using RunUMAP for visualization of nuclear transcriptomic heterogeneity. Cell types were manually annotated based on canonical marker gene expression. Differentially expressed genes (DEGs) were identified using the FindAllMarkers (Wilcoxon Rank Sum test) function in Seurat. Genes were further filtered by log_2_FC > 1 and adjusted P value < 0.05. Over-representation analysis (ORA) enrichment test was performed using the EnrichR function against the Gene ontology (GO) database. Gene Set Enrichment Analysis (GSEA) was completed using the ClusterProfiler R package. Enriched pathways were considered significant at an adjusted P value < 0.05.

### 19. Statistical Analysis

All data are presented as mean ± standard deviation (SD) unless otherwise specified. Unpaired two-tailed t-test was used for comparison of two groups. For analyses involving more than two groups, one-way or two-way ANOVA was performed, followed by Tukey’s multiple comparison test. Statistical analyses were conducted using GraphPad Prism (versions 8 or 10), and p < 0.05 was considered statistically significant. Significance levels were denoted as p < 0.05 (*), p < 0.01 (**), p < 0.001 (***), and p < 0.0001 (****). Images of tissue sections were taken from 3 locations (proximal, middle, distal) for each animal (at least n = 3). Images were analyzed using ImageJ (National Institutes of Health, Bethesda, MD, USA) for 5 randomly selected fields of each image.

## Acknowledgements

This work was supported by a grant from the National Institutes of Health, R01AG068250 to S.T.A. Schematics were created using Biorender (publication and licensing rights agreement MB24JBPCQV).

---

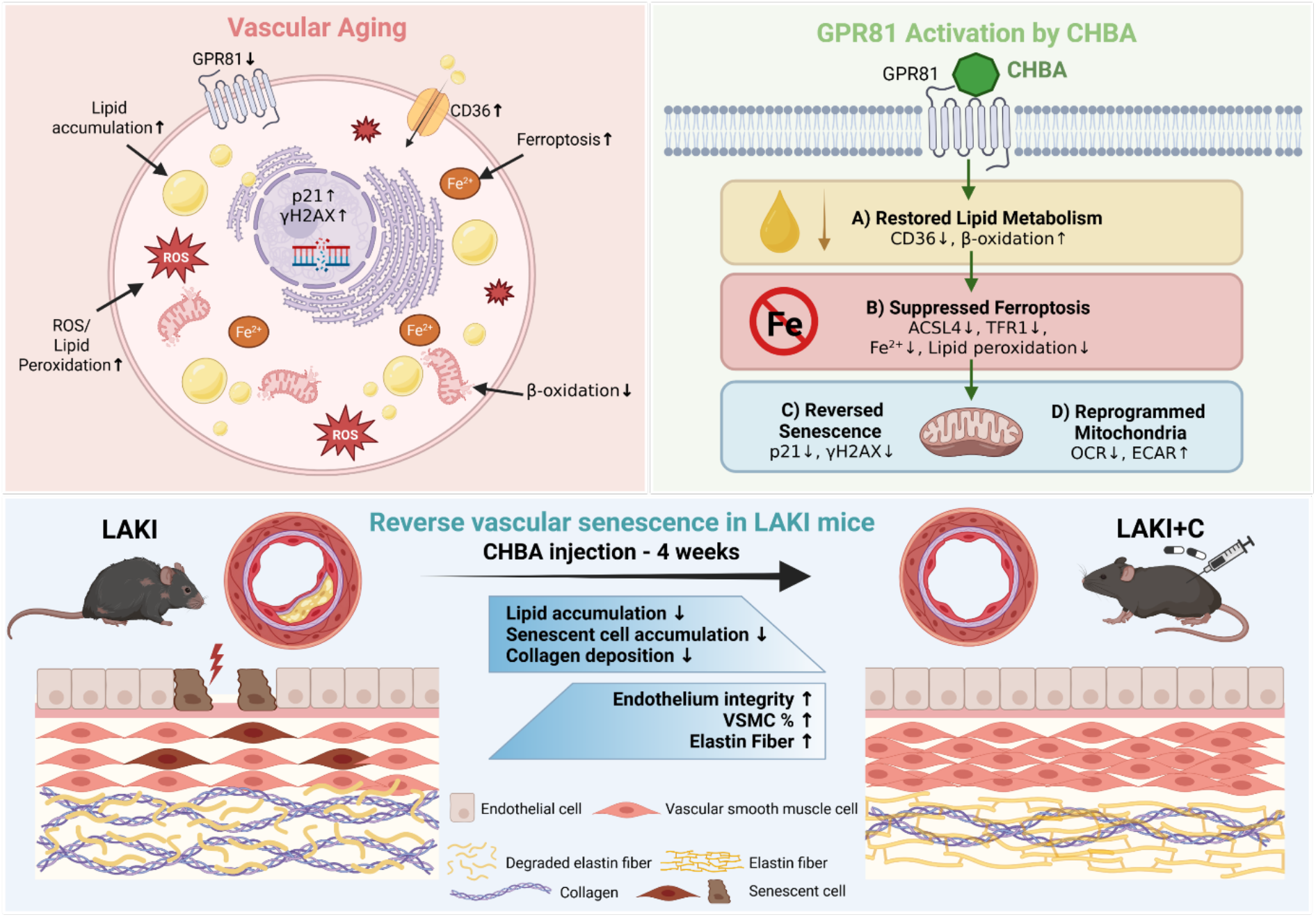

**Supplementary Figure S1:**
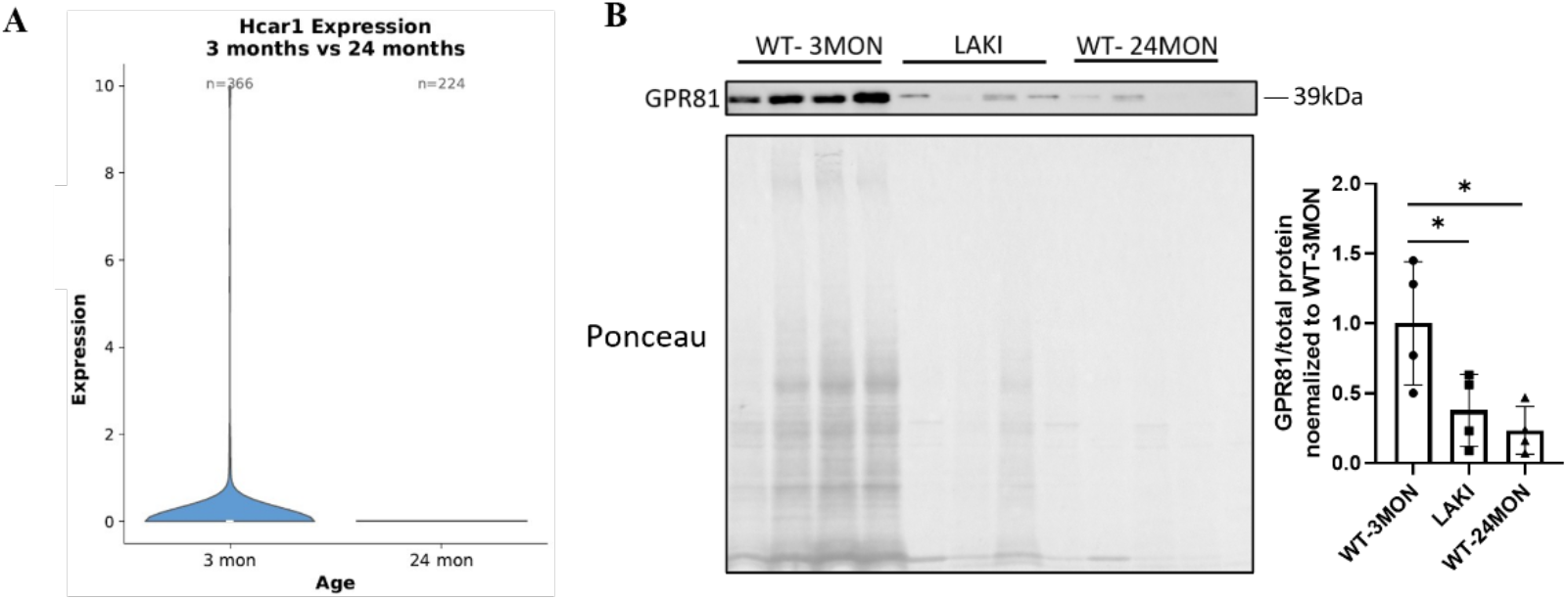
**A)** Violin plot showing Hcar1 expression in aortic cells from single-cell RNA-sequencing data of mouse aortas from 3-month-old and 24-month-old mice in the Tabula Muris Senis dataset. **B)** Representative western blot of GPR81 expression in aortas from 3-month-old wild type (WT), 24-month-old WT and LAKI mice.

**Supplementary Figure S2:**
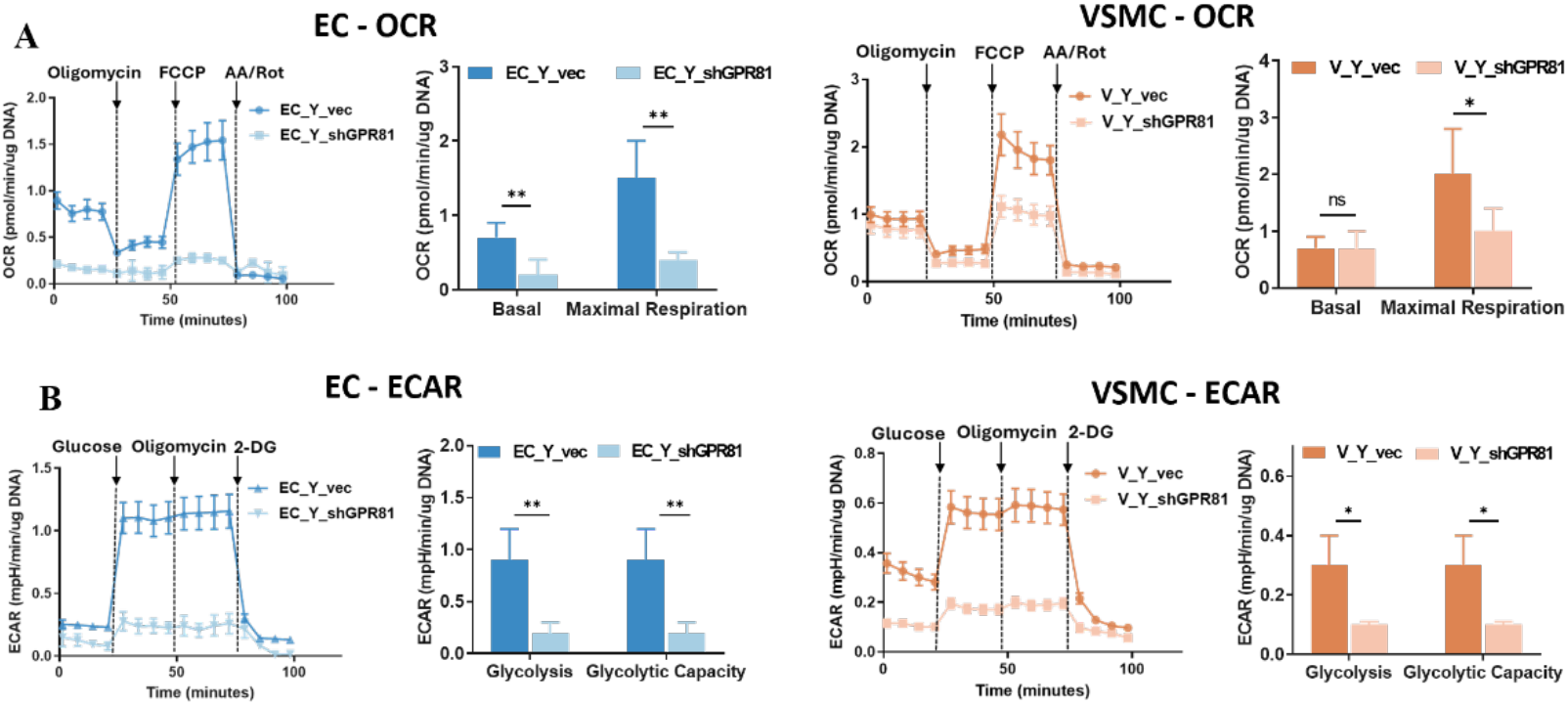
**A)** Seahorse extracellular flux analysis of mitochondrial respiration in EC (left, blue) and VSMC (right, orange), showing OCR traces following sequential injections (oligomycin, FCCP, antimycin A/rotenone) and quantification of basal and maximal respiration. Data shown as mean ± SD. **B)** ECAR measurements in EC (left) and VSMC (right) following sequential injections (glucose, oligomycin, and 2-DG), and quantification of glycolysis and glycolytic capacity. Data shown as mean ± SD.

**Supplementary Figure S3:**
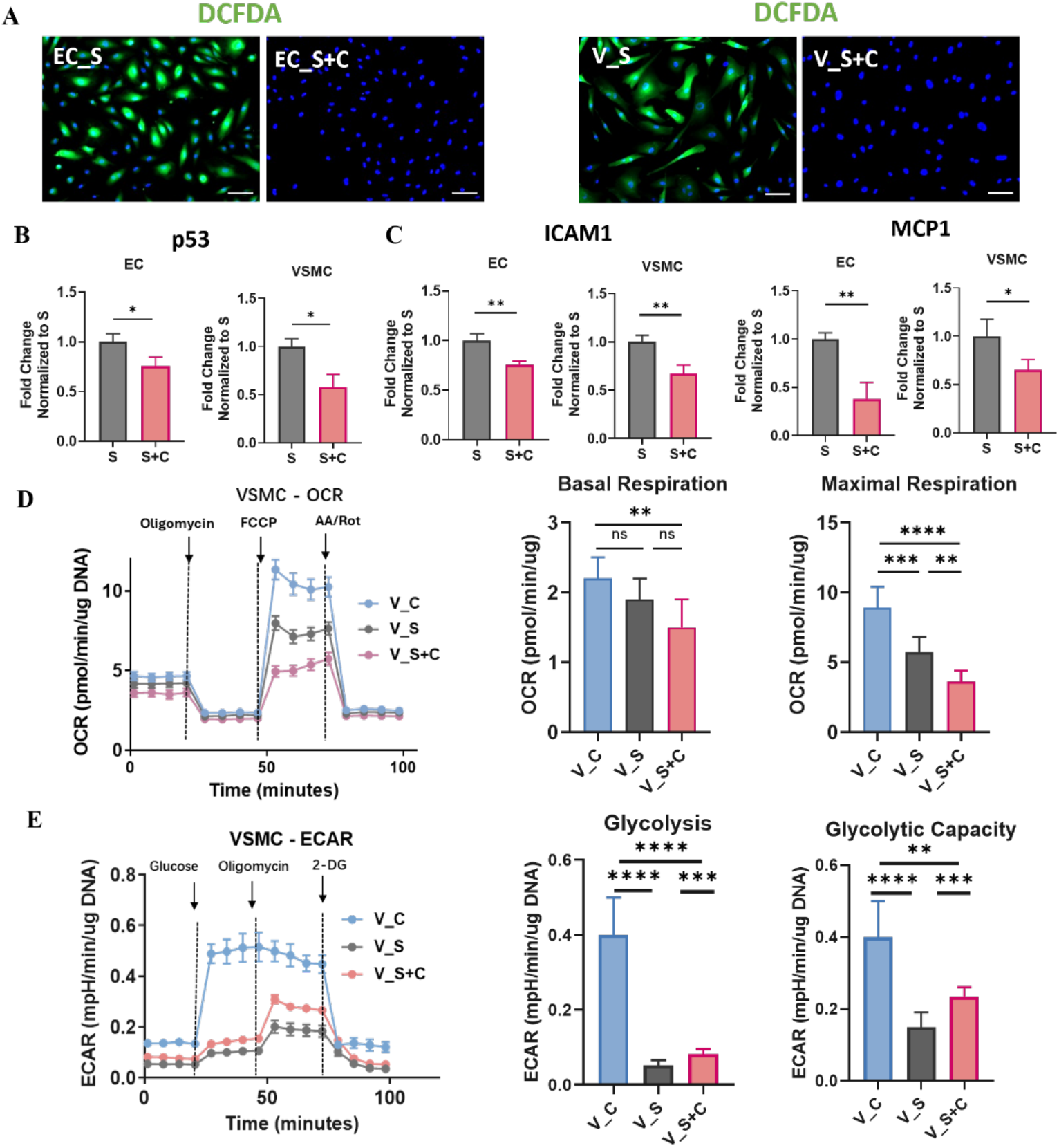
**A)** DCFDA staining to detect ROS in senescent ECs and VSMCs cultured without (S) or with CHBA (S+C). Scale bar, 100 µm. **B-C)** Quantitative RT-PCR for **(B)** p53, **(C)** ICAM-1 and MCP1. Data shown as mean ± SD. **D)** Seahorse extracellular flux analysis of mitochondrial respiration in VSMC, showing OCR traces following sequential injections (oligomycin, FCCP, antimycin A/rotenone); and quantification of basal and maximal respiration. Data shown as mean ± SD. **E)** ECAR measurements following sequential injections (glucose, oligomycin, and 2-DG), and quantification of glycolysis and glycolytic capacity. Data shown as mean ± SD.

**Supplementary Figure S4:**
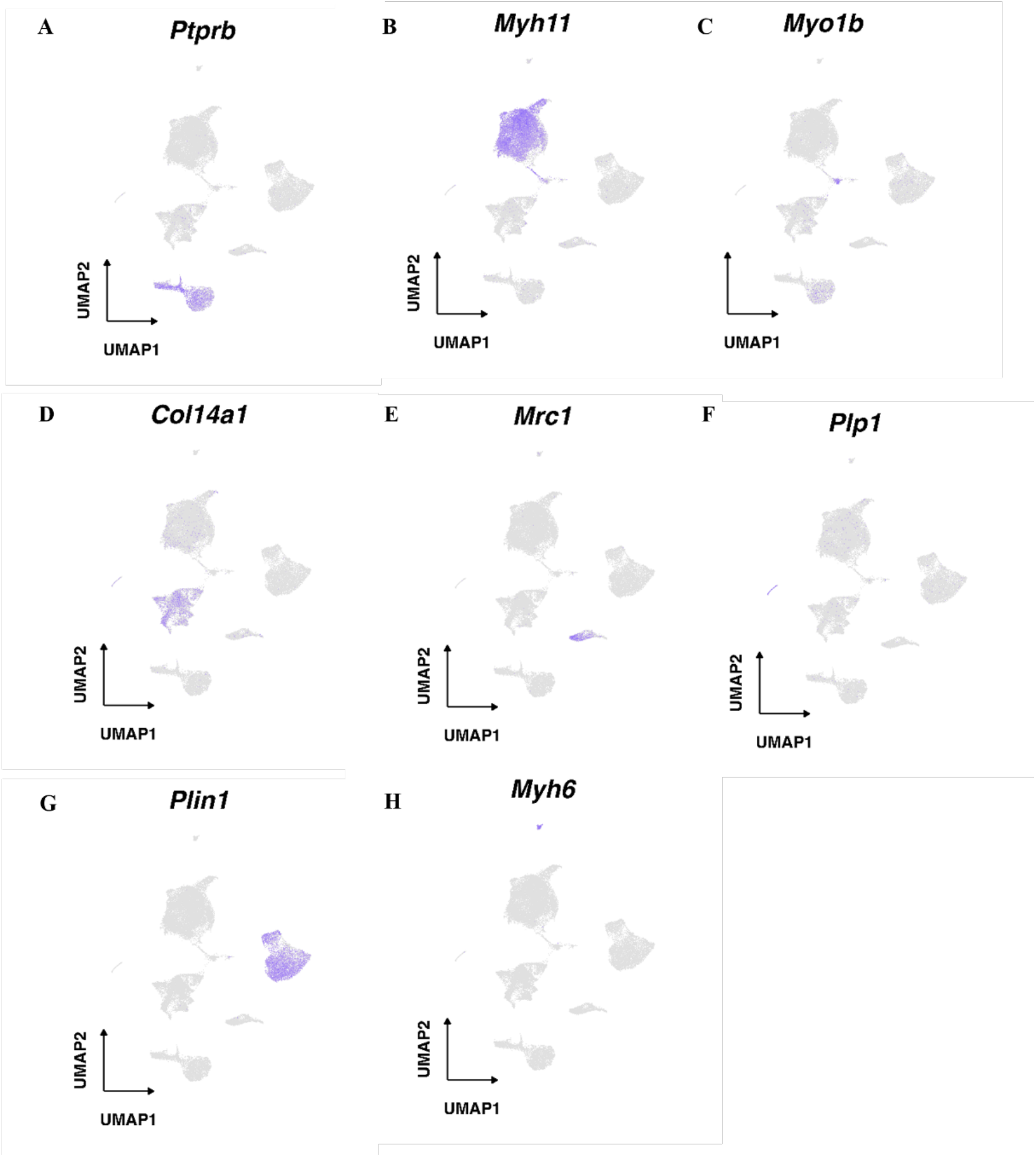
Gene expression feature plot showing expression of canonical markers used to define different cell clusters: **A)** Ptprb for ECs, **B)** Myh11 for VSMCs, **C)** Myo1b for Pericytes, **D)** Col14a1 for Fibroblasts, **E)** Mrc1 for Immune cells, **F)** Plp1 for Neural cells, **G)** Plin1 for Adipocytes, **H)** Myh6 for Myh6+ contractile cells.

**Supplementary Figure S5:**
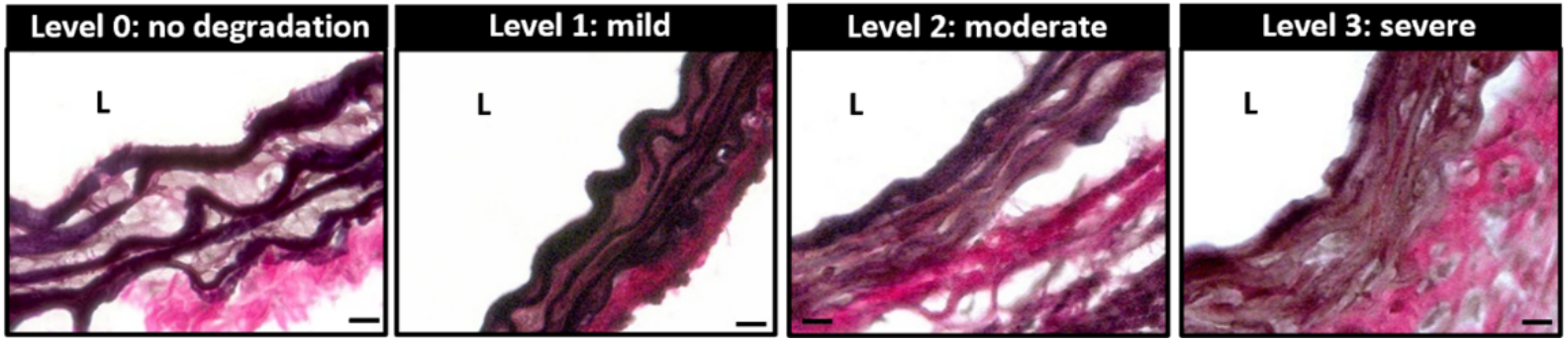
Representative images of Verhoeff staining in cross sections of arteries showing examples of each elastin degradation score. Scale bar represents 20 µm. L represents lumen.

**Supplementary Figure S6:**
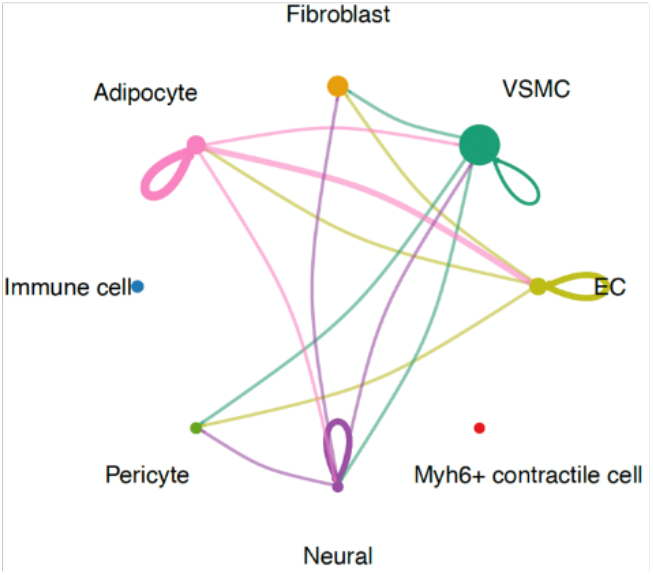
CellChat analysis showing the number of pairwise interactions among all cell clusters in WT arteries. Each node represents cell type, with node size proportional to the total number of interactions associated with that population. Connecting lines represent predicted ligand–receptor-mediated communication between cell types. Line color corresponds to the sender cell population and line thickness indicates the relative strength of communication.

**Supplementary Table 1:**
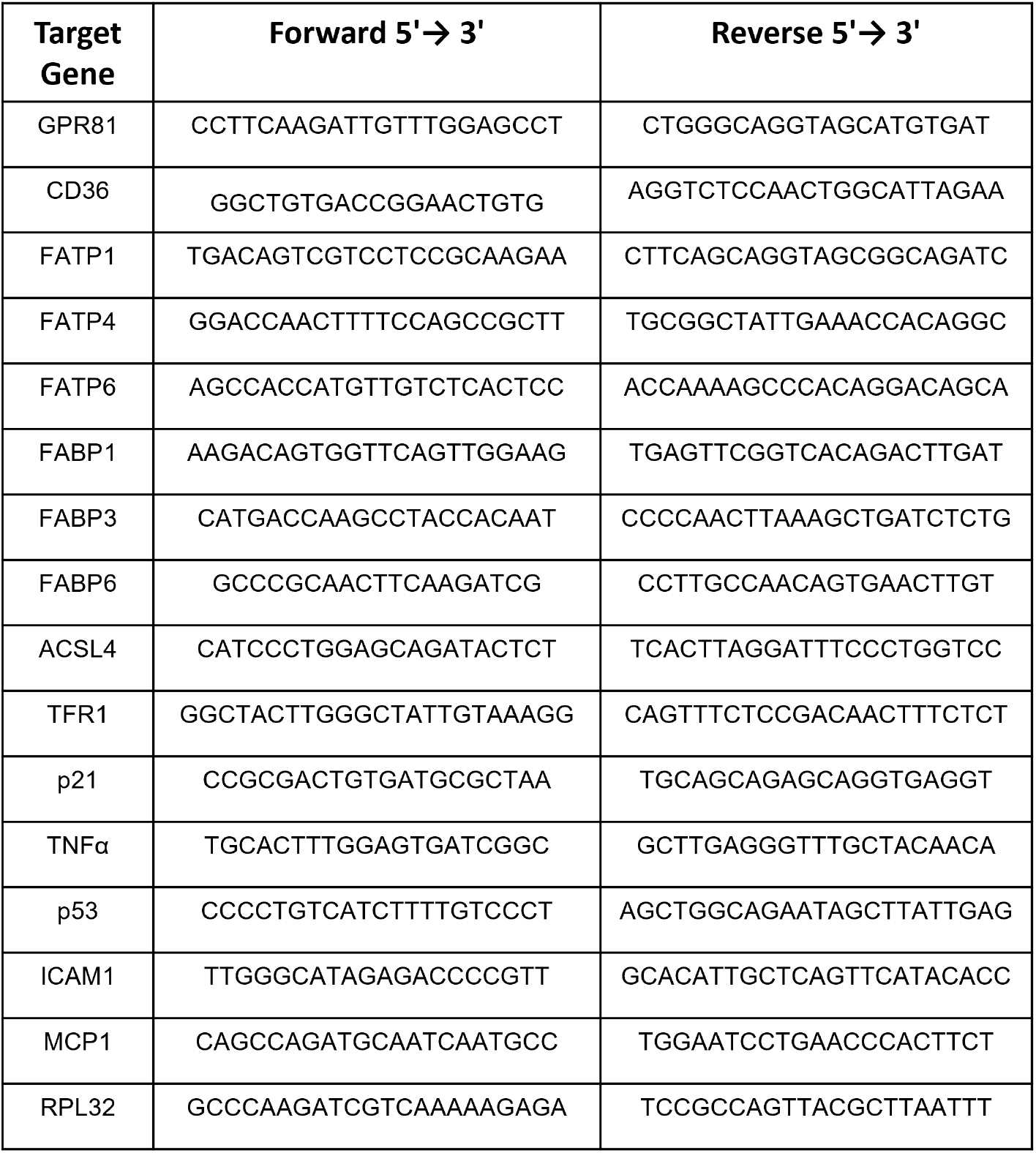
List of Primers.

**Supplementary Table 2:**
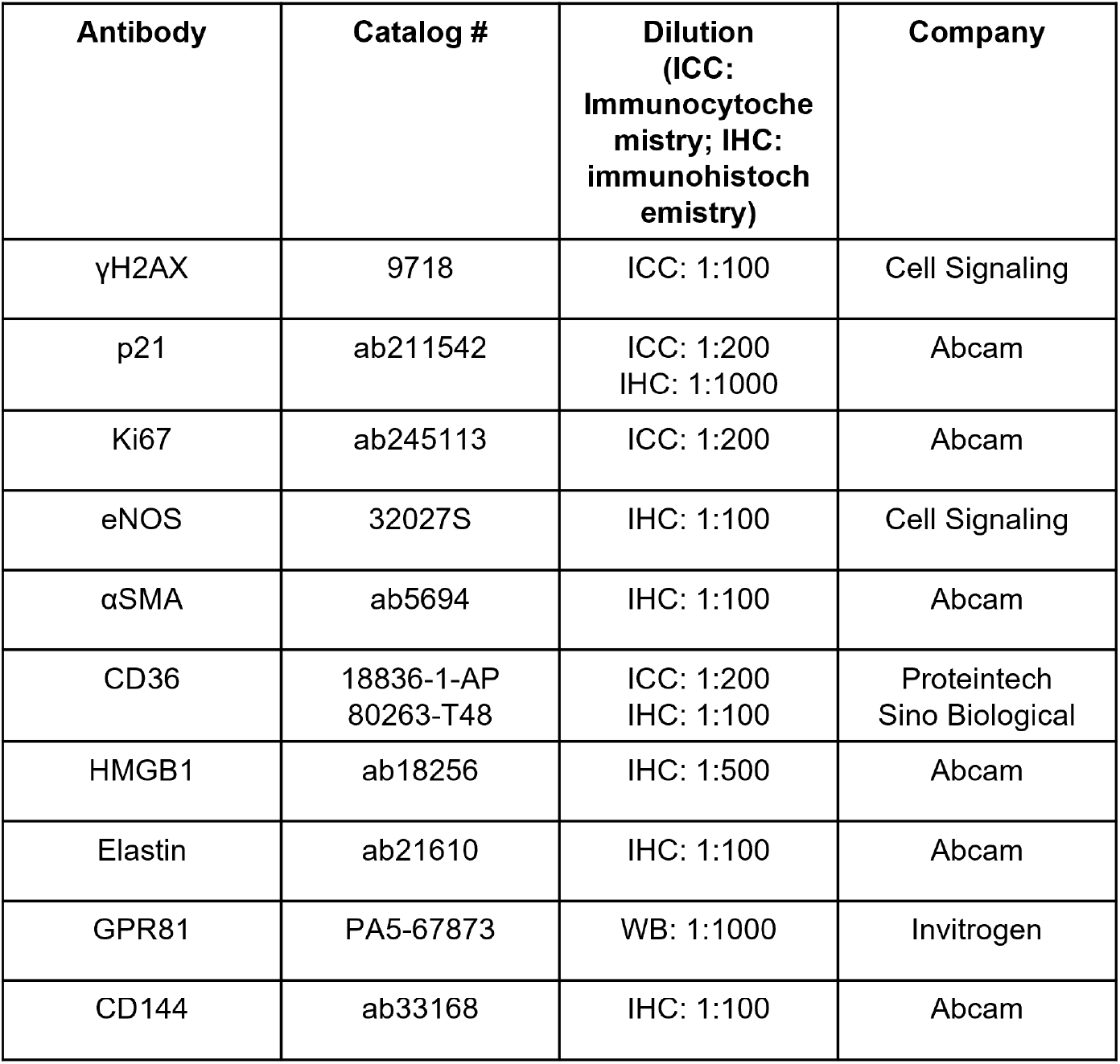
List of primary antibodies used in IHC/ICC.

| **Antibody** | **Catalog #** | **Dilution**<br>**(ICC:**<br>**Immunocytoche**<br>**mistry; IHC:**<br>**immunohistoch**<br>**emistry)** | **Company** |
| --- | --- | --- | --- |
| γH2AX | 9718 | ICC: 1:100 | Cell Signaling |
| p21 | ab211542 | ICC: 1:200<br>IHC: 1:1000 | Abcam |
| Ki67 | ab245113 | ICC: 1:200 | Abcam |
| eNOS | 32027S | IHC: 1:100 | Cell Signaling |
| αSMA | ab5694 | IHC: 1:100 | Abcam |
| CD36 | 18836-1-AP<br>80263-T48 | ICC: 1:200<br>IHC: 1:100 | Proteintech<br>Sino Biological |
| HMGB1 | ab18256 | IHC: 1:500 | Abcam |
| Elastin | ab21610 | IHC: 1:100 | Abcam |
| GPR81 | PA5-67873 | WB: 1:1000 | Invitrogen |
| CD144 | ab33168 | IHC: 1:100 | Abcam |

